# Rhizobiales commensal bacteria promote *Arabidopsis thaliana* root growth via host sulfated peptide pathway

**DOI:** 10.1101/2021.05.25.444716

**Authors:** Jana Hucklenbroich, Tamara Gigolashvili, Anna Koprivova, Philipp Spohr, Mahnaz Nezamivand Chegini, Gunnar W. Klau, Stanislav Kopriva, Ryohei Thomas Nakano

## Abstract

Root-associated commensal bacteria that belong to the order Rhizobiales, which also contains symbiotic and pathogenic bacteria, promote primary root growth of *Arabidopsis thaliana.* However, the molecular mechanism underlying this root growth promotion (RGP) activity remained unclear. Here, we conducted a transcriptomic analysis of *A. thaliana* roots inoculated with root-associated commensal bacteria of Rhizobiales and sister lineages and revealed common and strain/lineage-specific transcriptional response, possibly mediated by WRKY and ANAC family of transcription factors. We showed that the observed common response was also partly triggered by a wide range of non-pathogenic bacteria, fungi, and a multikingdom synthetic community (SynCom). This response was characterized by a down-regulation of genes related to intracellular redox regulation, suggesting distinctive redox status between pathogenic and non-pathogenic interactions. By integrating with developmental and cell biological experiments, we identified a crucial role of TYROSYLPROTEIN SULFOTRANSFERASE (TPST) in Rhizobiales RGP. Conversely, none of the known TPST-dependent sulfated peptide pathways appeared to be required for this activity, suggesting an unidentified component in the protein sulfation pathway targeted by Rhizobiales RGP. Finally, we show that TPST is needed for RGP exerted by Rhizobiales but not Pseudomonadales isolates, delineating lineage-specific mechanisms to manipulate host root development.

## Introduction

Various prokaryotic and eukaryotic microorganisms heavily colonize plant tissues in nature, collectively shaping the plant microbiota. Many of these do not engage in pathogenic or mutually symbiotic lifestyles and are therefore frequently referred to as “commensal” microbes (Berg et al., 2020). Bacterial communities associated with roots (root bacterial microbiota) are significantly distinct from soil microbiota (Bulgarelli et al., 2012; Lundberg et al., 2013) and qualitatively highly conserved across different plant species at the family level (Hacquard et al., 2015; Thiergart et al., 2020). However, at the quantitative and strain levels, there appears to be host species/genotype specificity (Wagner et al., 2016; Zgadzaj et al., 2016; Wippel et al., 2021; Yu et al., 2021). These findings indicate a regulatory mechanism in plants and bacteria that enables taxonomically defined community assembly in a host-dependent manner. It also implies that, throughout evolution, the extant microbiota members have benefited the host plants and *vice versa.* Recent studies have provided evidence that the root bacterial microbiota as a community is vital for plant health, for example, by protecting the host from fungi (Wei et al., 2015; Duran et al., 2018) or by facilitating nutrient acquisition (Castrillo et al., 2017; Harbort et al., 2020; Salas-Gonzalez et al., 2021). Imbalance of the microbiota compositions (“dysbiosis”) in roots as well as in leaves results in growth penalty (Chen et al., 2020; Finkel et al., 2020), illustrating the importance of eubiotic homeostasis of root microbiota (Paasch and He, 2021). Moreover, root bacterial microbiota members can interfere with root development and immunity. For example, many, in not all, commensal bacteria isolated from healthy *A. thaliana* roots can either promote or inhibit host primary root growth in mono-association (Finkel et al., 2020). Another subset of root commensals have been shown to suppress root growth inhibition triggered by a chronic exposure to flg22, a 22-amino acid immunogenic peptide derived from bacterial flagella (thereby referred to as suppressive isolates; Gomez-Gomez et al., 1999; Garrido-Oter et al., 2018; Yu et al., 2019; Ma et al., 2021; Teixeira et al., 2021). Overall, these studies have demonstrated the holobiome concept, an idea of collectively considering plants and associated microbiota as a single physiological and ecological unit. In contrast, our knowledge about the physiology of plants, including the developmental regulation of roots, has been primarily provided by studies using axenic plants. Toward an ultimate understanding of the plant molecular physiology under ecological conditions, it remains crucial to better describe the molecular dialog between plants and commensal bacteria and how this influences host physiological pathways.

Comparative analysis of root microbiota across a wide range of plant species has identified bacterial lineages highly prevalent across and abundant within root microbiota (Hacquard et al., 2015). These include Rhizobiales, a bacterial order within the class Alphaproteobacteria that contains strains engaging in different lifestyles. For example, rhizobia, the legume symbionts, induce nodule organogenesis in legume roots and fix atmospheric nitrogen into ammonium for the host growth, while Agrobacterium, a sublineage within the genus *Rhizobium*, pathogenically invade plant tissues to induce tumors and ultimately restrict plant growth. These lifestyles are highly differentiated and require dedicated genes in bacteria and plants, such as *nod/nif* and *NOD FACTOR RECEPTOR (NFR)* genes in rhizobia and legumes, respectively, and *vir* genes in Agrobacterium. In contrast, the vast majority of bacteria in the order Rhizobiales, including strains isolated from non-leguminous plants, animals, and soils, do not possess these genes (Garrido-Oter et al., 2018), and host-associated and soil free-living lifestyles are frequently interchangeable in this bacterial order (Wang et al., 2020). Therefore, commensals strains in the order Rhizobiales likely interact with host plants in a manner distinct from symbiotic and pathogenic lifestyles. We previously showed that the extant commensal Rhizobiales bacteria are commonly capable of promoting primary root growth of *Arabidopsis thaliana* Col-0 wild-type plants when gnotobiotically inoculated with germ-free hosts. Bacterial isolates from the sister lineages, Sphingomonadales and Caulobacterales, did not possess this host root growth promotion (RGP) activity, implying that the RGP activity represents an ancestral trait specific to the order Rhizobiales (Garrido-Oter et al., 2018). The wide prevalence among extant Rhizobiales strains also suggests an important role of the RGP activity in the interaction between *A. thaliana* roots and Rhizobiales commensals. However, the molecular mechanisms by which Rhizobiales bacteria interfere with host root growth remained unknown.

In the previous study, we showed that Rhizobiales RGP was associated with an enlargement of the root meristematic zone (MZ), where cells undergo proliferation without elongation or differentiation. On the other hand, the cell size in the elongation zone (EZ) and the differentiation zone (DZ) was comparable between axenic and R129_E-inoculated roots (Garrido-Oter et al., 2018). In *A. thaliana,* an orchestrated operation of various pathways regulates root cell growth and proliferation: The mitotic activity of stem cell daughters within the MZ is maintained by a gradient accumulation of PLETHORA (PLT) transcription factors (TFs) under the control of auxin (Aida et al., 2004; Galinha et al., 2007). Auxin accumulated within the MZ induces expression of *TYROSYLPROTEIN SULFOTRANSFERASE (TPST),* which encodes an enzyme to catalyze protein sulfation of proteins (Komori et al., 2009; Zhou et al., 2010), including ROOT MERISTEM GROWTH FACTOR (RGFs) (Matsuzaki et al., 2010; Shinohara, 2021). Recent studies showed that the perception of RGFs leads to an activation of MITOGEN-ACTIVATED PROTEIN (MAP) kinases and redistribution of reactive oxygen species (ROS) to increase the PLT protein accumulation (Lu et al., 2020; Shao et al., 2020; Yamada et al., 2020). The accumulation level of auxin and PLT gradually decreases along with the longitudinal axis of the roots, and an auxin minimum is formed at the transition zone (TZ), where cytokinin antagonizes with auxin to switch cell fate from division to elongation (Dello Ioio et al., 2008; Di Mambro et al., 2017). This transition process is also influenced by brassinosteroid and ROS distribution (Tsukagoshi et al., 2010; Chaiwanon and Wang, 2015). After the cell fate transition, the root cells continue growing within the EZ until they reach the DZ. This elongation in the EZ is facilitated by gibberellin and another group of sulfated peptides, PHYTOSULFOKINE (PSK) and PLANT PEPTIDE CONTAINING SULFATED TYROSINE 1 (PSY1) (Matsubayashi et al., 2006; Amano et al., 2007; Achard et al., 2009; Ubeda-Tomas et al., 2009; Ladwig et al., 2015), whose sulfation is also dependent on TPST (Komori et al., 2009). The *tpst* mutant of *A. thaliana* lacks all functional sulfated peptides, including but not limited to RGFs, PSKs, and PSY1, as well as another class of sulfated peptides involved in endodermal cell differentiation (CASPARIAN STRIP INTEGRITY FACTOR, CIFs; Doblas et al., 2017), and shows a highly disorganized root MZ, leading to severe root growth defects (Komori et al., 2009). It is plausible that Rhizobiales commensal bacteria target one of the processes mentioned above to exert the RGP activity. Our previous study showed that genetic perturbation of auxin-cytokinin antagonism in *arr1arr12* and *ahk3-3* mutants (Dello Ioio et al., 2008) did not disrupt Rhizobiales RGP (Garrido-Oter et al., 2018), pointing to a possible role of other components than auxin and cytokinin.

In this study, to disentangle the molecular interactions between *A. thaliana* and commensal Rhizobiales, we examined the host transcriptional response in roots inoculated with Rhizobiales and sister lineage commensal isolates. We identified common transcriptional responses commonly triggered by these bacteria. We then revealed that a part of the observed common response was also triggered by a wide range of non-pathogenic bacterial and fungal microbes. By analyzing lineage-specific responses and the developmental details of roots inoculated with Rhizobiales, we show that TPST, but none of the known receptors for RGF and PSKs, is indispensable for Rhizobiales RGP. Lastly, we show that many Rhizobiales but not Pseudomonadales commensals require TPST to promote root growth, illuminating the lineage-specific innovations of mechanisms to interfere with the host root development.

## Results

### Host transcriptional reprogramming by root-associated Alphaproteobacteria commensals

To obtain an overview of transcriptional responses to Rhizobiales commensals, we isolated and sequenced RNA from *A. thaliana* (Col-0 wild type) roots inoculated with a panel of root-associated commensals that belong to the class Alphaproteobacteria. Our previous RNAseq analysis revealed that the transcriptional responses triggered by R129_E were similar in roots at 4, 8, and 12 days post-inoculation (dpi), which were largely different from roots at 16 dpi (Garrido-Oter et al., 2018). A statistically insignificant or marginal RGP phenotype is typically observed at 12 dpi at the level of primary root length, while developmental manipulation at the cellular level (i.e. enlarged root MZ) already initiates at an earlier time point (at 4-8 dpi). Therefore, we decided to harvest roots at 12 dpi for our RNA-seq experiment to consider the impact of Rhizobiales RGP at the cellular level while minimizing the difference accounted for by different root growth. As inoculant bacteria, we used R129_E, R13_A, Root491, and Root142 from the order Rhizobiales and Root1497 and Root700 from the sister lineages Sphingomonadales and Caulobacterales, respectively, as outgroup controls. Among these isolates, only Rhizobiales strains exhibit root growth promotion (RGP) activity, while only R129_E and Root1497 are able to suppress host flg22-triggered root growth inhibition (Garrido-Oter et al., 2018). We identified a total of 1,439 differentially expressed genes (DEGs) between axenic and commensal-inoculated roots, based on the false discovery rate (FDR; α = 0.05) and log_2_-scale fold changes (logFC; |logFC| > 1), and a principal coordinates analysis (PCoA) was performed using their zero-centered expression values (Figure 1A). This revealed that these commensals triggered a significant shift in the host transcriptional profiles (R^2^ = 0.662, *P* < 0.001, PERMANOVA) in a manner dependent on the taxonomic lineages they belonged to (filled *versus* open shapes; R^2^ = 0.457, *P* = 0.0045, pairwise PERMANOVA between Rhizobiales and sister lineage isolates). We revealed a number of DEGs specific to each inoculation, as well as DEGs commonly identified in more than one inoculation (Supplemental Figure 1). On the other hand, transcriptional responses to these commensal bacteria were qualitatively similar and showed a strong correlation (Figures 1B and 1C), indicating that the taxonomy-dependency of host transcriptional reprogramming, as shown in Figure 1A, is explained by their quantitative differences.

**Figure 1.**
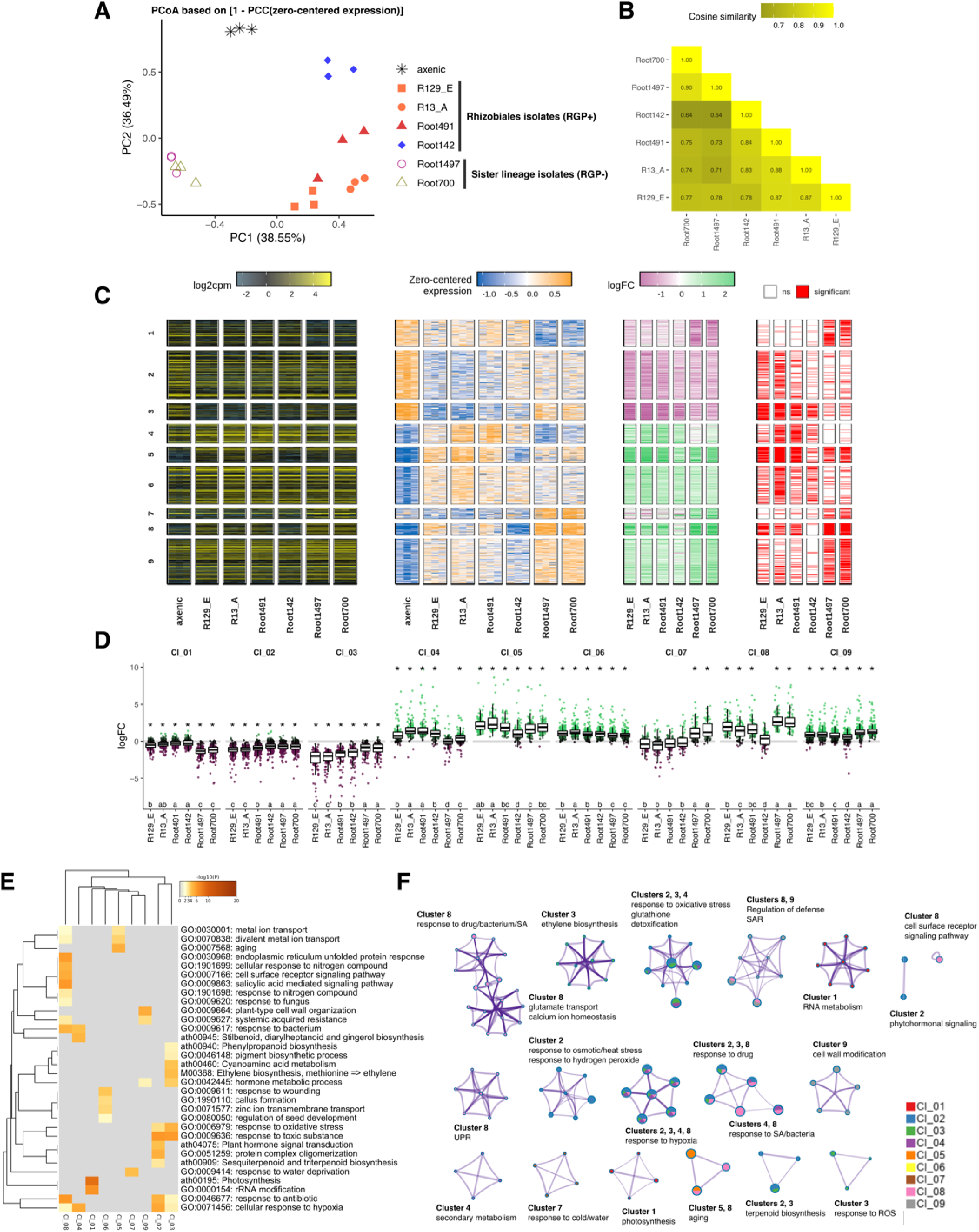
Alphaproteobacteria commensals trigger qualitatively similar transcriptional responses in host roots. (A) A principal coordinates analysis separated transcriptome of axenic roots (asterisks), roots inoculated with Rhizobiales commensals (closed shapes), and roots inoculated with sister lineage isolates (open shapes). (B) A heatmap showing cosine similarities between logFC upon Alphaproteobacteria commensal inoculations. (C) Heatmaps showing log_2_-scale count per million (log2cpm), zero-centered expression values, log_2_-scale fold changes (logFC), and significance (FDR < 0.05; ļlogFCļ > 1), aligned according to their k-means cluster assignment (*k* = 9, based on Bayesian information criterion). (D) Boxplots showing logFC of each *k*-means cluster. Asterisks indicate statistical significance corresponding to Student’s one-sample t test after Bonferroni’s correction for multiple comparisons (μ = 0; α = 0.001) and letters indicate statistical significance corresponding to pairwise Wilcoxon’s rank sum test after Bonferroni’s correction for multiple comparisons within each cluster (α = 0.05). (E) A heatmap showing enrichment of GO terms in the *k*-means clusters. (F) A network plot of enriched GO terms. Each node represents each GO term, and GOs with overlapping gene contents are connected with edges. *k*-means assignment of composing genes is shown as a pie chart. We selected the terms with the best *P* values from each of the 9 clusters, with the constraint that there are no more than 15 terms per cluster and no more than 250 terms in total. Representative terms are manually curated and indicated with composing *k*-means clusters for each GO cluster.

To obtain functional insights, we classified DEGs into 9 *k*-means gene clusters based on the zero-centered expression values and tested whether Gene Ontology (GO) categories were enriched in each cluster (Figures 1C to 1F and Supplemental Figure 2). We assigned down-regulated genes into Clusters 1, 2, and 3 and up-regulated genes into Clusters 4 to 9. Clusters 2 and 5 illustrated the general response to these commensal bacteria, and Clusters 3, 4, and 6, and Clusters 1, 7, and 9 represented responses quantitatively characteristic to Rhizobiales and sister lineage isolates, respectively. The commonly up-regulated Cluster 5 was enriched in genes related to metal ion transport and ageing, while commonly down-regulated Cluster 2 was enriched in genes related to response to hypoxia and to various abiotic stresses, as well as genes associated with biosynthesis of terpenoid, phytohormone signal transduction, and the detoxification pathway. Clusters characteristic to sister lineage isolates (Clusters 1, 7, and 9) were enriched in unique GO categories, such as photosynthesis (Cluster 1), response to cold and water deprivation (Cluster 7), and cell wall rearrangement and immune responses (Cluster 9). Clusters characteristic to Rhizobiales were characterized by up- and down-regulation of “Response to hypoxia” and “detoxification” (Clusters 3 and 4), which were also enriched in Clusters 2 as described above. Clusters 3, 4, and 6 were also characterized by down-regulation of biosynthesis of ethylene, glucosinolate, and flavonoid, and response to various abiotic stresses (Cluster 3), and up-regulation of response bacteria, wounding, salicylic acid (SA), and jasmonic acid (JA) (Clusters 4 and 6). Cluster 8 contained genes up-regulated by all isolates except for Root142, which were enriched in immune-related genes, such as genes responding to SA, bacteria, fungi, and hypoxia or related to calcium homeostasis, as well as genes involved in response to abiotic stresses such as heat, drugs, and amino acids. We previously reported that Root142 is one of the Rhizobiales strains colonizing *A. thaliana* roots efficiently (Garrido-Oter et al., 2018), implying the ability of Root142 to partly evade recognition by the host or to interfere with these host immune responses, albeit its inability to suppress flg22-triggered root growth inhibition. In contrast, only a handful (4 up- and 6 down-regulated genes) of DEGs were found to be common between suppressive R129_E and Root1497 (Supplemental Figure 1), probably accounted for by different mechanisms employed by these isolates to interfere with host immune responses. Nevertheless, these findings illustrate that the tested Alphaproteobacteria commensals commonly interfere with the host genetic pathways associated with hypoxia, detoxification, and plant immunity, to a different extent depending on the isolates. Lineage-specific DEGs (46 and 99 genes specific to Rhizobiales and the sister lineage, respectively) and isolate-specific DEGs (from 44 to 164 genes) also demonstrated the presence of lineage/strain-specific responses, whose functional aspects remained unclear based on their GO annotation.

We then looked for TF-binding motifs enriched in the 1-kb upstream sequences of the genes in each cluster. We identified several binding motifs, mainly from the TF families of WRKY and ANAC, as well as the AHL family of DNA-binding proteins (Figure 2A), suggesting a role of these TF families in commensal-triggered transcriptional reprogramming. We found that 20 WRKY and NAC family TFs were differentially expressed (differentially expressed TFs, DETFs) and used these DETFs as a list of potential regulators to infer a gene regulatory network (GRN) by the so-called GENIE3 algorithm (Figure 2B). This algorithm considers both positive and negative potential regulations to infer a GRN, which is a strong advantage over correlation-based approaches, such as *k*-means clustering and gene co-expression networks (Huynh-Thu et al., 2010). The inferred GRN was composed of two arbitrarily defined modules, characterized by either WRKY or NAC family DETFs as hub nodes (module 1 and module 2, respectively; Figure 2B). Among DETFs, WRKY54 and ANAC060 showed the first and second highest centrality indices (degree, closeness, and betweenness centralities), suggesting their role in the observed transcriptional reprogramming. Genes commonly induced/suppressed by Alphaproteobacteria commensals (Clusters 1, 8, and 9) were enriched in module 1, whereas genes in the Rhizobiales-characteristic clusters (Clusters 3, 4, ad 6) were enriched in module 2 (Supplemental Figures 3A and 3B). We observed similar trends using all DETFs, independent of promoter motif enrichment analysis, as potential regulators to infer a GRN, albeit with a less clear GRN structure (Supplemental Figure 4). Importantly, in both GRNs, WRKY54 and ANAC060 were identified as candidate hub regulators based on the centrality indices and connected with genes whose transcriptional responses were commonly triggered by all isolates or characteristically by Rhizobiales isolates, respectively. Together, our RNAseq experiments demonstrated common and lineage-specific transcriptional responses in roots that were potentially mediated by WRKY54 and ANAC060, respectively, along with other TFs.

**Figure 2.**
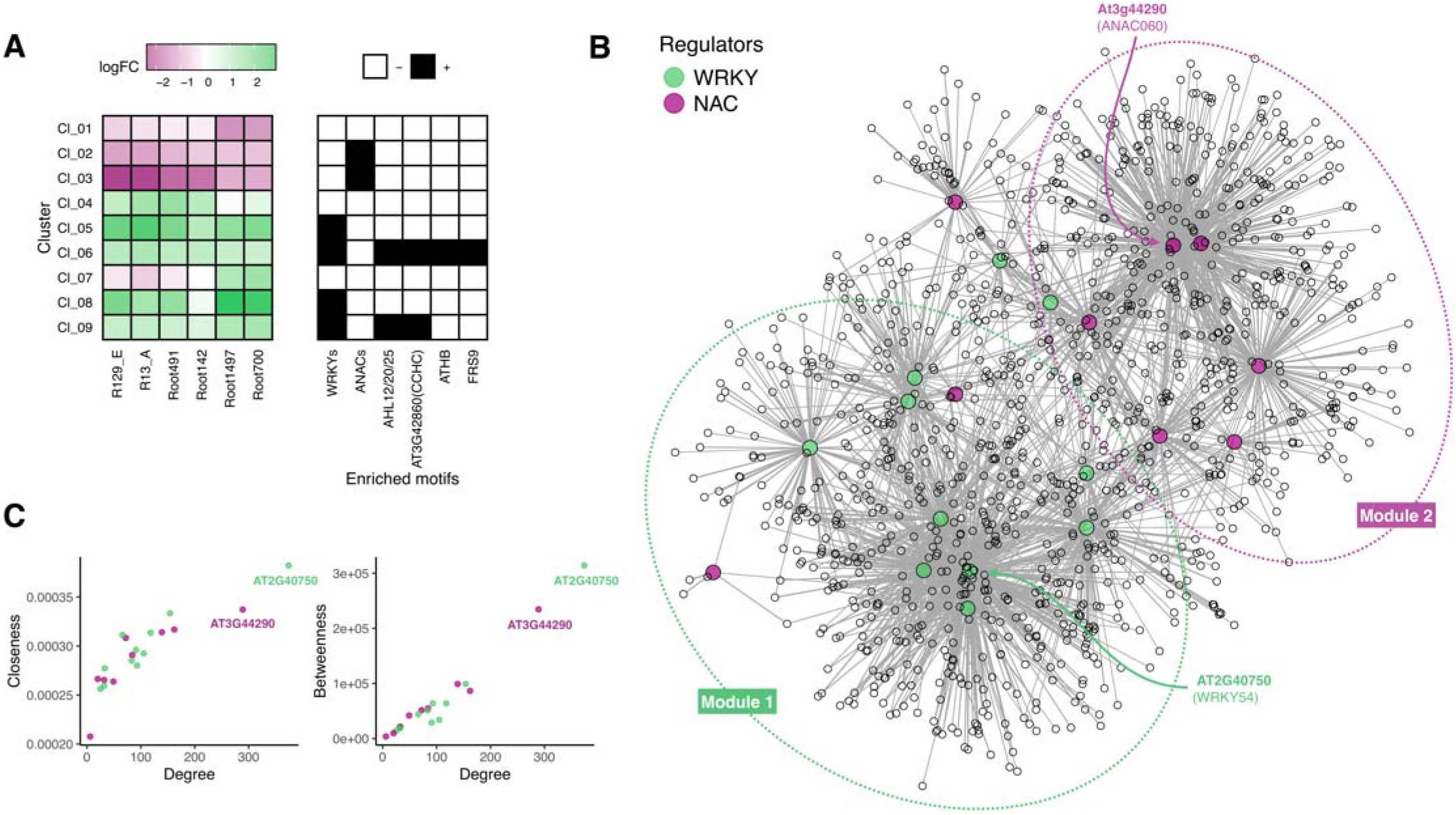
WRKY and ANAC transcription factors are important for transcriptional reprogramming. (A) Heatmaps showing mean logFC of each cluster (left) and the enrichment of corresponding transcription factor binding motifs in their promoter regions (1-kb upstream from transcription initiation sites). AHL, AT-HOOK MOTIF CONTAINING NUCLEAR LOCALIZED; CCHC, CCHC-type zinc-finger motif containing protein; ATHB, ARABIDOPSIS THALIANA HOMEOBOX; FRS9, FAR1-RELATED SEQUENCE9. (B) GENIE3-inferred gene regulatory network using WRKY (green) and ANAC (magenta) family DETFs as potential regulators (colored; bigger node size). Arbitrarily defined modules are shown by dotted lines. (C) Degree, closeness, and betweenness centralities were computed based on the network in (B) for all potential regulators. Green and magenta dots correspond to WRKY and ANAC family TFs.

### Overlapping transcriptional responses to a wide range of non-pathogenic microbes

Our RNAseq experiments identified common as well as lineage-specific transcriptional responses, both of which were enriched in similar GO terms, such as “detoxification” and “response to hypoxia”. We noted that these GO terms were also found among genes responding to other root microbiota members (Hacquard et al., 2016; Ma et al., 2021; Teixeira et al., 2021; Wippel et al., 2021). This suggests that a wide range of root-associated microbes might broadly interfere with this response, and we aimed to understand the extent to which the observed root transcriptional response is common and specific to Alphaproteobacteria and Rhizobiales commensals. We compared our data with publicly available transcriptomic datasets using the roots inoculated with root-inhabiting microbes that engage in beneficial, commensal, and pathogenic lifestyles under different nutrient conditions, in different experimental setups, at different time points (Supplemental Table 1). These include beneficial and pathogenic *Colletotrichum* fungi *(C. tofieldiae* and *C. incanum;* Hacquard et al., 2016; Hiruma et al., 2016), a bacterial synthetic community (B-SynCom; Harbort et al., 2020), and a multi-kingdom synthetic community (SynCom) composed of bacteria, fungi, and oomycetes (BFO-SynCom; Hou et al., 2020). In addition, we sequenced RNA obtained from roots inoculated with beneficial and pathogenic *Burkholderia* bacteria *(B. phytofirmans* and *B. glumae* isolated from onion and rice, respectively), as well as from roots inoculated with R129_E in the presence or absence of flg22 to consider the impact of host immune responses. We compared and clustered responses to different microbial inoculations based on the similarity of logFC (Figures 3A). LogFC within non-pathogenic inoculations was significantly more similar than between the responses to pathogenic and non-pathogenic inoculations (Figure 3B), without any clear sign of the taxonomy or the complexity of respective inocula (shapes in the dendrogram). A PCoA based on the transcriptome-wide logFC similarities revealed a clustering of non-pathogenic inoculations, irrespective of their taxonomy and complexity (Figure 3C). A pairwise PERMANOVA between different lifestyles (commensal, beneficial, pathogenic, and SynCom) revealed that all combinations of lifestyles significantly contribute to the difference in the host transcriptional responses (FDR-corrected *P* values of 0.002 ~ 0.01), while the R^2^ values were larger between pathogenic and non-pathogenic lifestyles than within non-pathogenic lifestyles (Supplemental Table 2). Notably, minimizing batch effects by a canonical analysis of principal coordinates (CAP) clustered all non-pathogenic interactions with microbes isolated from *A. thaliana* (Figure 3D).

**Figure 3.**
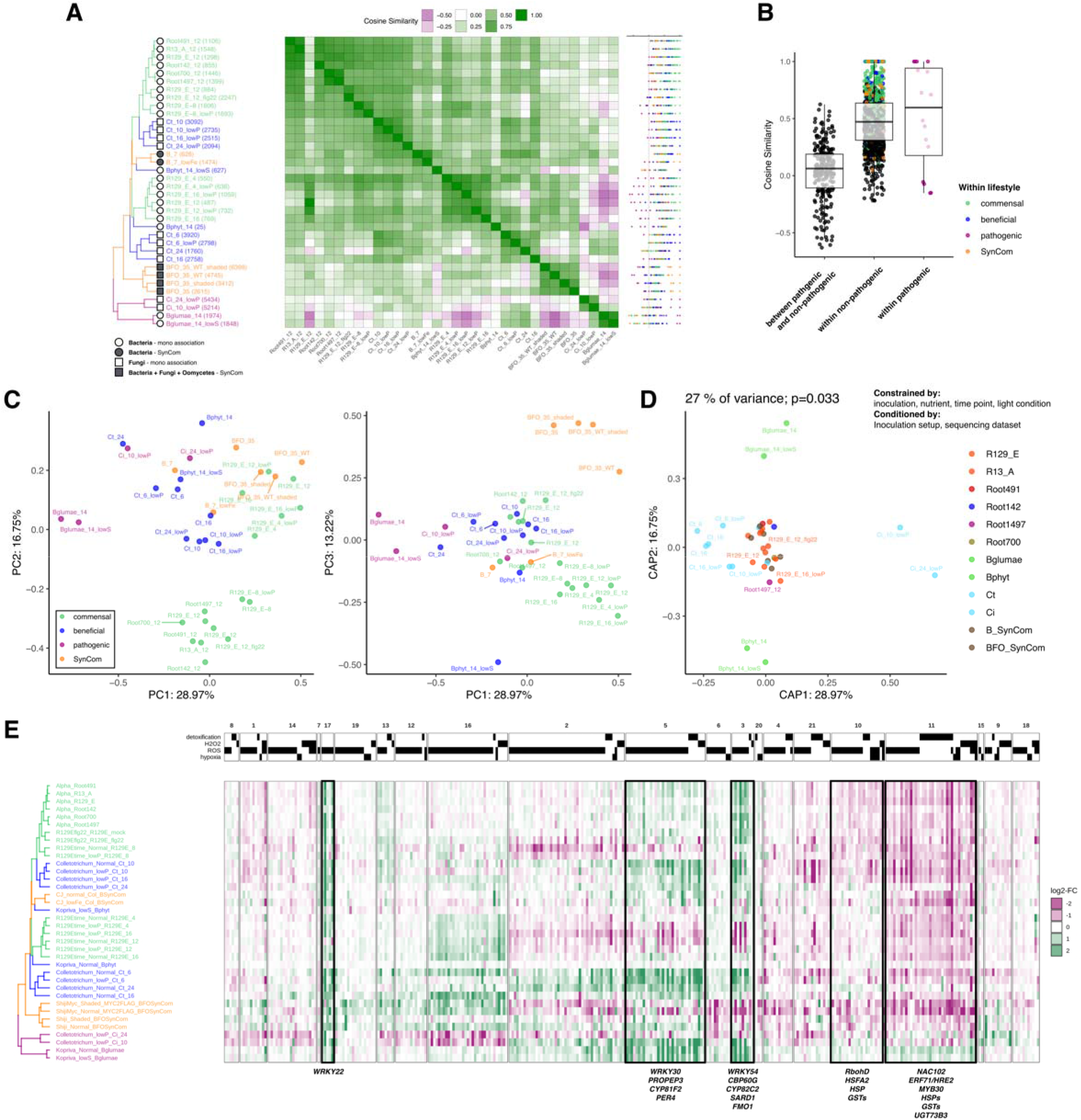
Non-pathogenic microbial inoculation triggered similar transcriptional responses in the host roots. (A) A heatmap showing cosine similarities between logFC upon different microbial inoculations. DEGs were selected based on the inoculation in rows (numbers of DEGs are shown in parentheses), whose logFC cosine similarity was calculated in comparison with inoculations in columns. For each row, all cosine similarity values were plotted on the right, whose colors indicate the lifestyle of the comparing inoculation. (B) Cosine similarities within non-pathogenic inoculations and within pathogenic inoculations were higher than those between pathogenic and non-pathogenic inoculations. (C) Principal coordinates analysis based on logFC cosine similarities of all commonly quantified genes (n = 21,815). Colors correspond to lifestyles. (D) Constrained analysis of principal coordinates (CAP) based on the logFC cosine similarities, using the constraining and conditioning factors indicated above. Colors correspond to inoculated microbial strains. (E) A heatmap showing logFC of genes assigned with the GO categories related to detoxification and response to hydrogen peroxide, reactive oxygen species, and hypoxia. Inoculations are aligned according to the dendrogram in (A) and genes are aligned according to k-means clustering as shown in Supplemental Figure 2B. Numbers indicated in the names of inoculation indicate the time point of harvesting (dpi).

Concatenated lists of DEGs (13,889 genes in total) were then classified into 21 *k*-means clusters and tested for their enriched GO categories (Supplemental Figures 5A and 5B). This identified the detoxification pathway to be strongly enriched in Cluster 11 (*P* = 5.1 ×10^-18^), whose transcriptional suppression characterized the non-pathogenic interactions. Our of 22 genes in Cluster 11 assigned with the GO term “detoxification”, 13 genes encode glutathione (GSH) *S*-transferases (GSTs), which play an important role in cellular redox homeostasis. The hypoxia response is known to trigger ROS production via RESPIRATORY BURST OXIDASE HOMOLOG D (RbohD) in *A. thaliana* (Pucciariello et al., 2012), and is associated with strong immune response induction. Therefore, we speculated that non-pathogenic interactions may impact the root redox response. Among DEGs, we found a total of 327 genes assigned with GO categories of response to hypoxia, response to ROS, and detoxification pathway (Figure 3E). We found that the hypoxia-inducible immune-related genes, such as *FLAVIN-DEPENDENT MONOOXYGENASE 1 (FMO1;* Hartmann et al., 2018), *CALMODULIN-BINDING PROTEIN 60-LIKE G (CBP60G;* Wang et al., 2009), *ELICITOR PEPTIDE 3 PRECURSOR (PROPEP3;* Huffaker and Ryan, 2007), *CYTOCHROME P450 FAMILY 82, SUBFAMILY C, POLYPEPTIDE 2 (CYP82C2;* Rajniak et al., 2015), *PEROXIDASE4 (PER4;* Rasul et al., 2012), and *WRKY22* (Hsu et al., 2013), were mostly up-regulated in many pathogenic and non-pathogenic interactions (Clusters 3/5/17), suggesting that the genetic sector responding to commensal bacteria inoculation overlaps with the hypoxia response. On the other hand, another subset of hypoxia-responsive genes, such as *RbohD, HEAT SHOCK TRANSCRIPTION FACTOR A 2 (HSFA2), ARABIDOPSIS NAC DOMAIN CONTAINING PROTEIN 102 (ANAC102),* and *HYPOXIA RESPONSIVE ERF2 (HRE2),* along with many *GST* genes, were found in Cluster 11 and down-regulated by non-pathogenic microbes. Down-regulation of the transcriptional levels of *GSTs* was not limited to a subset of isoforms but broadly observed across most of the *GST* genes encoded by the *A. thaliana* genome (Supplemental Figure 5C), suggesting a link to the general redox response rather than a specific secondary metabolic pathway (discussed below). Overall, these results suggest that the response to hypoxia and the detoxification pathway, linked to intracellular root redox status, is associated with the interactions with non-pathogenic microbes, rather than representing the Rhizobiales- or Alphaproteobacteria-specific responses.

### ANAC060 transcription factor characterized Rhizobiales-specific root transcriptional response

Our initial RNAseq experiment identified the hypoxia response and detoxification pathway in the genes enriched in Rhizobiales-characteristic gene clusters (Clusters 3 and 4 in Figure 1), while the subsequent meta-analysis revealed that these gene pathways were rather associated with interaction with a wide range of microbes. Clusters 3, 11, and 17 contained more than 40% of Alphaproteobacteria DEGs (Supplemental Figure 5A, rank abundance plot) and contained genes related to hypoxia-responsive immunity and detoxification pathways (Figure 3E), suggesting that the hypoxia response and detoxification pathways are not necessarily associated with the Rhizobiales RGP activity. In order to characterize the molecular mechanism responsible for Rhizobiales-triggered RGP, we aimed to further explore the response characteristic to Rhizobiales independently of *k*-means clustering and Gene Ontology annotations. In our initial RNAseq dataset with Alphaproteobacteria commensals, we identified a total of 46 genes that were significantly up- or down-regulated by Rhizobiales but not by sister lineage isolates (“Rhizobiales DEGs”; Figure 4A), which were then mapped onto the GRN shown in Figure 2B. We found that Rhizobiales DEGs were exclusively located on a sub-module within module 2 and tightly associated with *ANAC060* (Figures 4B and 4C and Supplemental Figure 3C). Given that *ANAC060* is one of the two DETFs with the highest centralities (Figure 3C) and the only TF among Rhizobiales DEGs (Figure 4A), our results suggest that ANAC060 is crucially involved in the Rhizobiales-specific transcriptional response, a part of which ultimately triggers RGP phenotype.

**Figure 4.**
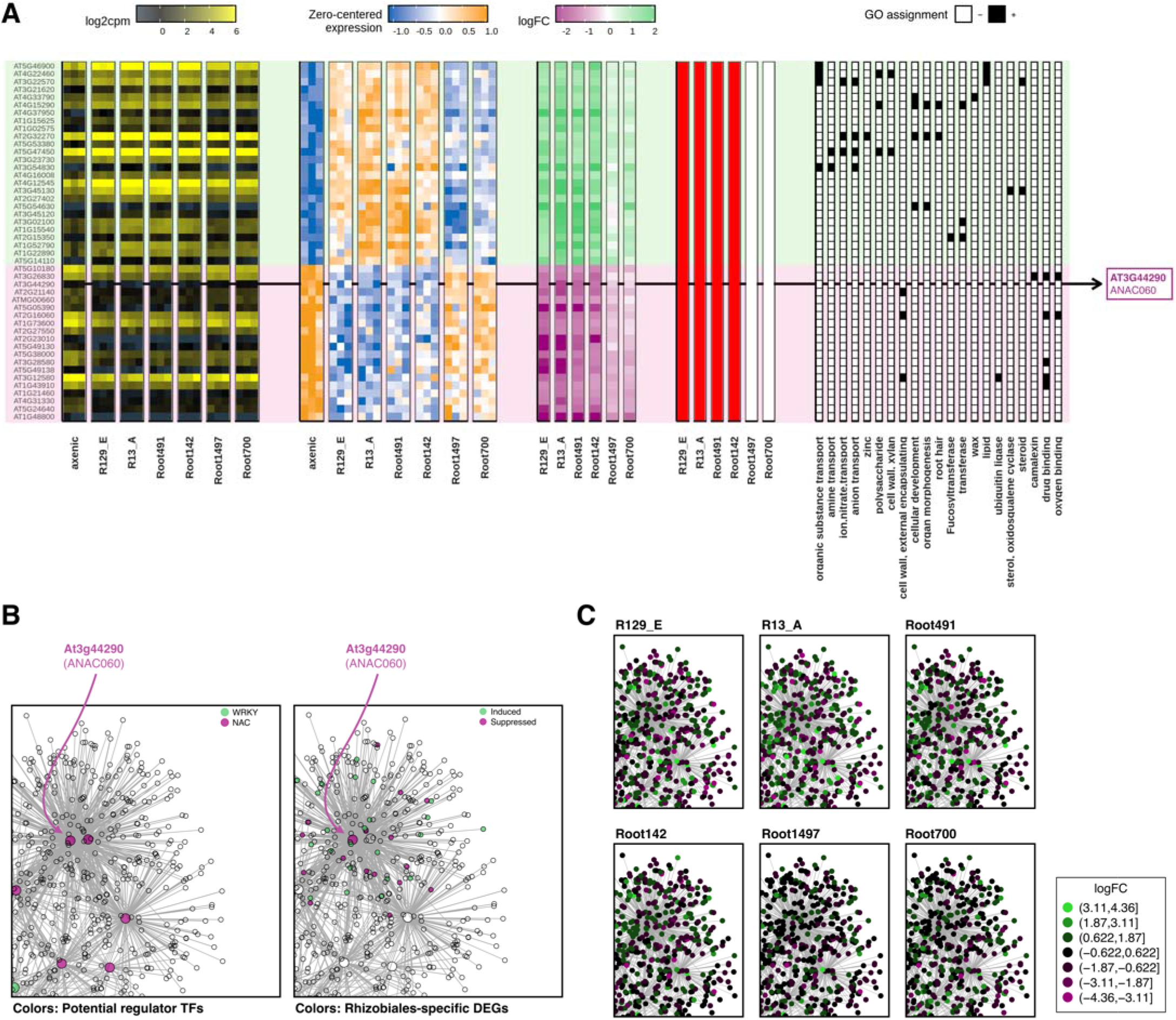
Rhizobiales-specific DEGs are potentially regulated by ANAC060. (A) Heatmaps showing log2cpm, zero-centered expression values, logFC, significance, and GO assignment of genes specifically regulated by Rhizobiales commensals but not by sister lineage isolates. (B) Enlarged view of the GENIE3-inferred GRN, shown in Figure 2B, colored for potential regulator TFs (left) or Rhizobiales-specific DEGs (right). (C) Same view as in (B), colored by their logFC upon different bacterial inoculation.

### RGP exerted by R129 E requires host sulfated peptide pathway

We previously reported that RGP in Rhizobiales is associated with an increase in the number of cells within the root MZ but not the cell size in EZ and DZ, pointing to facilitated cell division triggered by Rhizobiales commensals. We confirmed this by staining cells undergoing S-phase using a thymidine analog 5-ethynyl-2’-deoxyuridine (EdU; Supplemental Figure 7A), corroborating the scenario that Rhizobiales RGP targets root MZ homeostasis. On the other hand, we did not find a dramatic increase or decrease of auxin and cytokinin, at least not at the level of DR5:ER-GFP and TCSn:ER-GFP fluorescence intensity (Supplemental Figures 7B and 7C), which further supported the potential role of other components than auxin-cytokinin antagonism. In published single-cell RNA sequencing (scRNA-seq) data of *A. thaliana* roots (Denyer et al., 2019), *ANAC060* showed relatively high expression in the cell population characterized by the expression of meristematic genes (“cluster 11”; Supplemental Figures 6A and 6B). We also found that the top 100 genes co-expressed with *ANAC060* across 4,727 sequenced root cells were enriched in the GO categories related to root MZ regulation, such as “meristematic growth,” “regulation of cell division,” and “response to gravity” (Supplemental Figure 6C). Interestingly, genes that were best co-expressed with *ANAC060* in the scRNAseq dataset included *RGF3* (14th), as well as *PLT1* (5th) and *PLT2* (63rd) (Supplemental Figures S6A). Moreover, a pervious study reported that ANAC060 potentially represses the expression of PSY1 while activating the expression of PSY4, a sequence-related homologue of PSY1 (Yu et al., 2020; Tost et al., 2021), overall suggesting a potential link between ANAC060 and the sulfated peptide pathway.

To test the link between Rhizobiales RGP and the plant sulfated peptide pathway, we inoculated a Rhizobiales commensal R129_E with *tpst* mutant plants, depleted in all classes of functional sulfated peptides. Strikingly, we identified a lack of the RGP phenotype in the *tpst* mutant (Figure 5A), without a clear depletion of bacterial cells from roots (Figure 5B). In contrast, we observed significant RGP in *rgi1;2;3;4;5-1, pskr1pskr2psy1r,* and *sgn3* mutants, which are impaired in perception of RGF, PSK, and CIF peptides, respectively (Figure 5C and Supplemental Figure 8A). Likewise, the RGP activity of R129_E remained functional when the three major isoforms of RGF peptides operating in the root MZ homeostasis were genetically abolished in the *rgf123* mutant (Supplemental Figure 8A) (Matsuzaki et al., 2010). Disorganized root MZ in the *tpst* mutant is known to be rescued by external application of RGF1 peptide but not its desulfated form (dRGF1). However, treatments with neither of these peptides rescued the RGP activity of R129_E, despite the substantial recovery of root MZ under these conditions (Supplemental Figures 8B and 8C). These findings demonstrate that R129_E requires the host sulfotransferase enzyme TPST for its RGP activity but not the RGF and PSK peptide pathways. Of note, the root MZ in the *rgi1;2;3;4;5-1* mutant was even more severely disrupted than in the *tpst* mutant (Ou et al., 2016; Song et al., 2016), while *rgi1;2;3;4;5-1* nonetheless exhibited the RGP phenotype when inoculated with R129_E (Figure 5C). Combined with the observation that restoration of root MZ by RGF1 did not rescue RGP activity, our findings point to a specific regulatory role for sulfated peptides, making the potential confounding effects due to the pleiotropic effect of highly disorganized root MZ in *tpst* less likely. Lastly, we inoculated wild-type and *tpst* mutant plants expressing PLT1-GFP under its *cis*-regulatory sequence to test whether R129_E showed any impact on PLT1 protein accumulation. This showed that R129_E indeed stimulates the accumulation of PLT1-GFP in Col-0 root MZ, which was not observed when the sulfated peptides were abolished in *tpst* (Figure 5D). Overall, these findings indicate that the host sulfated peptide pathway is indispensable for R129_E-stimulated accumulation of PLT1 protein, which facilitates the cell division within the root MZ and contributes to promoting the primary root growth.

**Figure 5.**
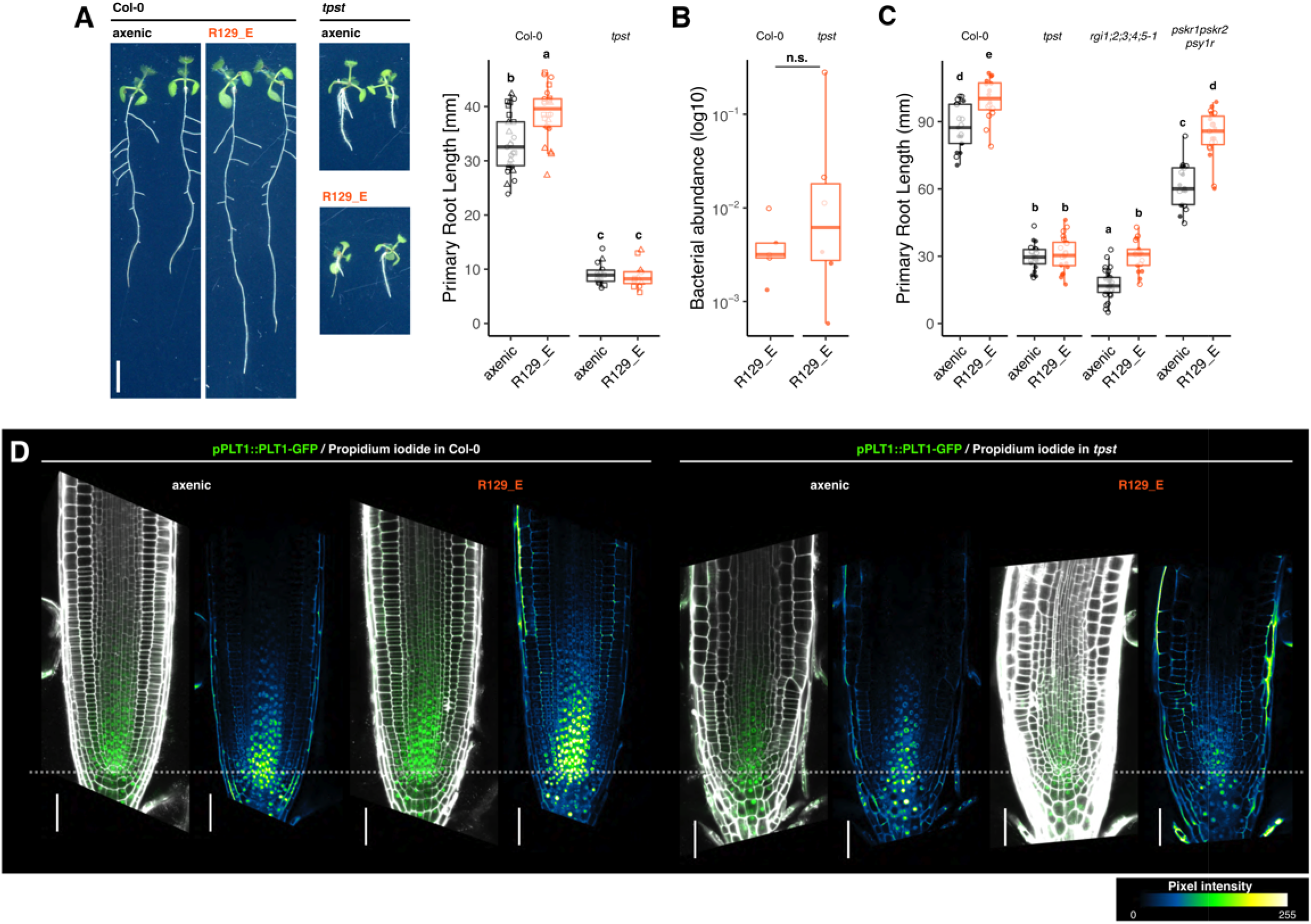
Plant sulfated peptide pathway is needed for R129_E RGP activity. (A) Primary root growth and representative pictures of Col-0 and *tpst* plants grown axenically or inoculated with R129_E. Seeds were inoculated and grown under continuous light conditions for 12 days. (B) Bacterial abundance relative to plant biomass was quantified by DNA-based quantitative PCR using primers specific to *A. thaliana* and R129_E *16S rRNA* genes. No clear depletion of bacterial cells from *tpst* roots was observed. (C) Mutants impaired RGF1 perception *(rgi1;2,3;4;5-1)* or PSK perception *(pskr1pskr2psy1r)* did not phenocopy *tpst* when 7-day-old seedlings were inoculated with R129_E for 14 days under short-day conditions. (D) Col-0 root MZ accumulate higher amounts of proPLT1::PLT1-GFP when 7-day-old seedlings were inoculated with R129_E for 7 days under short-day conditions, whereas *tpst* root MZ accumulated a similar amount irrespective of R129_E inoculation. For each panel, a merged image of propidium iodide staining (gray) and PLT1-GFP (green) signals and an image only showing PLT1-GFP signals pseudo-colored as shown in the calibration bar are shown in left and right, respectively. Bars indicate 50 μm. Images are aligned with quiescent centers on the dotted line. Letters indicate statistical significance (α = 0.05) corresponding to Student’s *t* test after fitting to a linear mixed model with biological and technical replicates as random factors, corrected for multiple comparison by the Benjamini-Hochberg method.

### The sulfated peptide pathway is needed by Rhizobiales but not Pseudomonadales RGP commensals

We previously showed that the RGP activity is prevalent within the order Rhizobiales (Garrido-Oter et al., 2018), while it remained unclear whether they promote root growth by the same molecular mechanism. We, therefore, inoculated *tpst* mutants with two additional Rhizobiales isolates, Root491 *(Agrobacterium)* and Root142 *(Sinorhizobium),* which were both used in the RNAseq analysis (Figure 1), and found that none of these isolates were able to promote *tpst* mutant root growth (Figure 6A). In contrast, when the *tpst* mutant was inoculated with Pseudomonadales isolates, namely Root7, Root71, Root329, and Root569, which are also capable of promoting *A. thaliana* root growth, we observed significant RGP phenotypes (Figure 6B). These findings indicate a specific and conserved contribution of the plant sulfated peptide pathway to Rhizobiales RGP, but not to Pseudomonadales RGP, illustrating the lineage-specific innovations of mechanisms to manipulate host root growth. This also implies that root-associated commensal bacteria have independently acquired the RGP capability more than once, pointing to a potential physiological relevance of this phenotype in the context of plantmicrobiota interactions.

**Figure 6.**
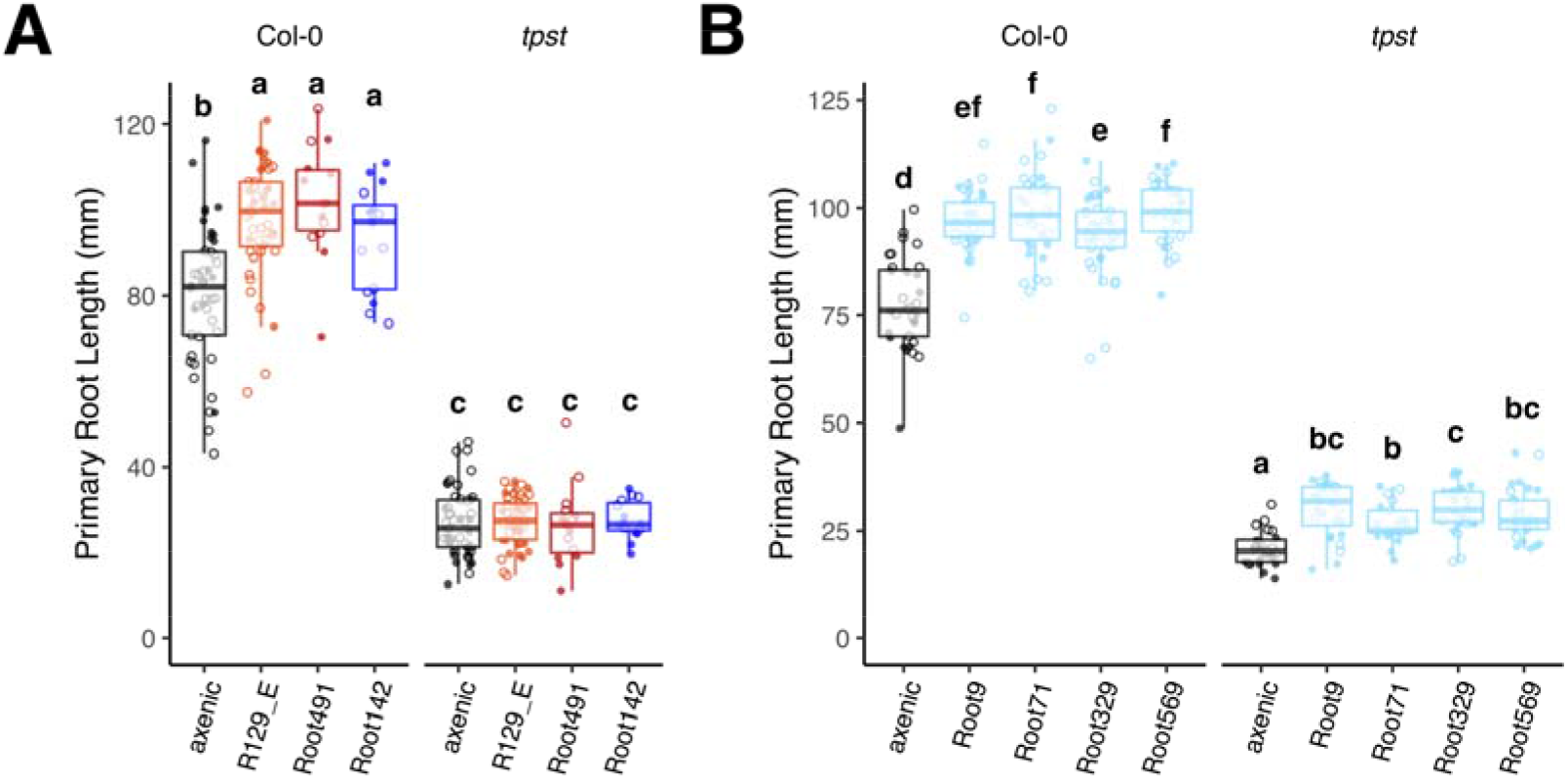
TPST is needed for the RGP activity explicitly exerted by Rhizobiales RGP but not Pseudomonadales. Seven-day-old seedling of Col-0 and *tpst* were inoculated with a panel of Rhizobiales commensals (A) or Pseudomonadales commensals (B) and grown for 14 days under short-day conditions. Letters indicate statistical significance (α = 0.05) corresponding to Student’s t test after fitting to a linear mixed model with biological and technical replicates as random factors, corrected for multiple comparison by the Benjamini-Hochberg method.

## Discussion

### WRKY and ANAC family TFs in transcriptional responses to Alphaproteobacteria commensals

Our comparative transcriptomic analysis identified genes that were differentially regulated by a panel of Alphaproteobacteria commensals. This enabled us to identify common as well as strain/lineage-specific responses, which were qualitatively similar and functionally overlapped with each other. The 1-kb upstream sequences of these genes tend to possess binding motifs of WRKY and ANAC family TFs, suggesting the role of these families of TFs in Alphaproteobacteria commensal-triggered transcriptional reprograming. WRKY family TFs are widely known for their crucial role in defense-related stress responses and are often involved in plant development, such as leaf senescence (Phukan et al., 2016). Many ANAC family TFs are reported to be involved in stress-induced regulation of development (Shao et al., 2015) and plant immunity (Yuan et al., 2019). It is well known that the WRKY family of TFs form a highly rigid transcriptional network (Eulgem and Somssich, 2007), and disruption of a single gene is frequently insufficient for perturbation of the entire network. This also appears to be the case for the ANAC family of TFs (Kim et al., 2018), pointing to a complex transcriptional network that ultimately contributes to the commensal-triggered reprogramming. The TF families identified in our promoter enrichment analysis were also found in a recent study addressing the co-occurrence of multiple TF binding motifs in promoters of the genes responding to immunogenic stimuli in roots in a cell type-specific manner (Rich-Griffin et al., 2020), further supporting the idea of their key role in plant-microbe interactions in roots of *A. thaliana.*

### Host redox status may play a role in the distinction between pathogenic and non-pathogenic microbes

Recent studies described transcriptional responses in roots and leaves to a different set of *A. thaliana* root- and leaf-derived bacterial commensals than those used in our study (Maier et al., 2021; Nobori et al., 2021; Teixeira et al., 2021). These commensal bacteria triggered qualitatively similar responses in the host, which is consistent with our finding that the response to Alphaproteobacteria commensals was largely similar. Furthermore, we have developed a method to compare transcriptional responses analyzed in independent RNAseq experiments. This allowed us to identify a substantial overlap between root transcriptional responses to a wide range of bacteria, fungi, and SynComs, inoculated in very different experimental setups, expanding the model of common transcriptional responses to a broader range of interactions. This includes up-regulation of hypoxia-responsive immune-related genes by both pathogenic and non-pathogenic interactions even after weeks of inoculation without causing growth restriction. This is contradictory with the growth-defense tradeoff model, according to which a chronic activation of immunity results in growth inhibition (Gomez-Gomez et al., 1999; Huot et al., 2014), suggesting the presence of an immune response sector that is uncoupled from the tradeoff. We also identified genes specifically and commonly down-regulated by non-pathogenic microbes, which were enriched in the detoxification pathway. We noted that this was mostly driven by the down-regulation of genes encoding GSTs. GSTs are often involved in secondary metabolic pathways, for example, by conjugation of GSH to toxic intermediates (thereby annotated as detoxification pathway components). Moreover, GSH is one of the critical components in sensing and regulating the cellular redox status, where GSTs play a crucial role (Noctor et al., 2012). We observed that the down-regulation of GSTs was not limited to GST isoforms that were specific to the family Brassicaceae (Supplemental Figure 5D, shaded by blue), which are often involved in lineage-specific secondary metabolism (Dixon et al., 2010), but were broadly observed among different classes of GSTs, including GSTUs and GSTFs (Pislewska-Bednarek et al., 2018). These results point to a possibility that pathogenic and non-pathogenic microbes differentially influence host cellular redox status. The broad impact on the expression of multiple GST genes also implies that their transcriptional down-regulation may be a consequence of altered GSH homeostasis, rather than a cause of the transcriptional reprogramming. A recent study suggested that the extracellular and cytoplasmic ROS responses are mutually distinguishable, where the cytoplasmic response is more prolonged (Arnaud et al., 2021). GST proteins in *A. thaliana* do not have predicted signal peptides for secretion (based on the prediction by SignalP-5.0) and are mainly localized in the intracellular compartments (Dixon et al., 2009; Noctor et al., 2012; Lallement et al., 2014). Given that we observed the influence of non-pathogenic inoculations on genes related to cellular redox at relatively late time points (from 4 days to 5 weeks), it is plausible that the cytoplasmic redox regulation is involved in the non-pathogenic interactions, while apoplastic ROS production is involved in the defense response against pathogens. Overall, based on these findings, we speculate that host root intracellular redox status has a role in the discrimination between pathogenic and non-pathogenic interactions, either as a determinant of the response or as a consequence of different types of interactions. Yet, the key molecular signatures in plants and microbes that differentiate pathogenic and non-pathogenic interactions remain to be identified.

### Rhizobiales commensals interfere with the host sulfated peptide pathway to promote root growth

We here provide clear genetic evidence for the role of plant sulfated peptides in Rhizobiales RGP. Plant sulfated peptides characterized thus far play a crucial role in plant development, including cell division, growth, and differentiation (Kaufmann and Sauter, 2019). In roots, RGFs, PSKs, and PSY1 orchestrate with a phytohormonal network to coordinate cell division and growth constitutively as well as in response to environmental conditions (Matsubayashi et al., 2006; Amano et al., 2007; Matsuzaki et al., 2010; Cederholm and Benfey, 2015), while CIFs specifically regulate endodermal cell differentiation (Doblas et al., 2017). Our mutant analysis demonstrated that known receptors of these sulfated peptides were not essential for Rhizobiales RGP. This suggests the presence of unidentified cognate receptors or yet another class of sulfated peptides. Receptors for extracellular peptides are typically a member of the leucine-rich repeat receptor-like kinase (LRR-RLK) superfamily, which has been immensely diversified in land plant lineages (Shiu et al., 2004; Furumizu et al., 2021). To date, only a handful of LRR-RLKs have been functionally characterized, while the physiological function of many members, including their cognate ligands, remains unknown. Of note, while the *pskr1pskr2psy1r* mutant is insensitive to PSKs, it retains the ability to respond to the PSY1 peptide, albeit to a lesser extent than wild-type plants (Pruitt et al., 2017). Given that the direct binding between PSY1 and PSY1R has not yet been experimentally shown, it is sensible to assume the presence of other LRR-RLKs that perceive PSY1 and related peptides (Tost et al., 2021). Besides, a large variety of uncharacterized secreted peptides are encoded by the *A. thaliana* genome, including several classes of peptides with a predicted motif for tyrosyl sulfation (aspartate-tyrosin; DY motif; Ghorbani et al., 2015), pointing to an unexplored repertoire of plant sulfated peptides. Further characterization of the mechanisms by which Rhizobiales promote root growth in a manner dependent on the host sulfated peptide pathway may lead to identifying a novel class of receptors and/or ligand sulfated peptides.

Interestingly, a hemibiotrophic bacterial pathogen *Xanthomonas oryzae* pv. *oryzae (Xoo)* also produces a sulfated peptide, whose amino acid sequence is highly similar to that of plant PSY1. This peptide, called RaxX, is needed for *Xoo* to exert its virulence in rice, which is, on the other hand, recognized by the host immune receptor XA21 to initiate responses that limit *Xoo* infection (Pruitt et al., 2015; Pruitt et al., 2017). Exogenous treatment of *A. thaliana* plants with RaxX promotes root growth, as does PSY1, and, therefore, it has been speculated that RaxX interferes with the PSY1 pathway to interrupt host immunity. Although RaxX has been the only bacterial sulfated peptides that have been molecularly characterized thus far, it is plausible that other bacterial species, especially those interacting with eukaryotic hosts, have acquired the ability to secrete sulfated peptides that mimic and/or interfere with host extracellular peptide signaling. In fact, genes with high similarity to the sulfotransferase RaxST in *Xoo* were found in *A. thaliana* root-associated bacterial genomes (20 out 194 genomes, based on blastp with an *E* value cutoff of 1.0; (Bai et al., 2015), mainly in those belonging to the orders Sphingomonadales, Burkholderiales, Pseudomonadales, and Xanthomonadales. It is also possible that even despite the lack of sequence-related RaxST homologs, other bacterial species have convergently acquired another class of enzymes that catalyze the sulfation of secreted peptides. Yet, it is not highly likely that bacterial sulfated peptides mediate the Rhizobiales RGP. It is important to note that the TPST enzyme catalyzes tyrosine *O*-sulfation within the Golgi apparatus, most likely at the *cis*-side, before the substrate (pro)peptides reach the apoplast after its translation in the endoplasmic reticulum (Komori et al., 2009). For external peptides to be sulfated by the host TPST, they need to be internalized into the host cells and transported to *cis*-Golgi, followed by tyrosine sulfation by TPST and re-secretion back to the apoplast. Overall, it is probable that the lack of TPST in the host does not directly affect *de novo* production of sulfated peptides by bacterial or mediate sulfation of naïve bacterial peptides in the extracellular space. Instead, our results delineate the involvement of plant endogenous sulfated peptides, which are yet to be identified in the following study.

### Physiological relevance of Rhizobiales RGP

Manipulation of host root growth appears to be a common feature of root-associated commensal bacteria (Zamioudis et al., 2013; Finkel et al., 2020), while one of the most common and dominant pathways is auxin-triggered root growth inhibition (Finkel et al., 2020). Despite the known mechanisms to manipulate host abiotic stress responses via the phytohormonal network (Bulgarelli et al., 2013), little is known about the mechanisms by which root commensal promote root growth. Moreover, it is yet to be seen whether root growth phenotypes observed in the artificial gnotobiotic system are relevant in the ecological/physiological context. Due to the plasticity and high variability of root growth depending on the environmental conditions, it remains technically challenging to directly tackle this question using wild-type plants. Here, we provide genetic evidence for the crucial role of sulfated peptides in Rhizobiales RGP. Retained RGP exerted by Pseudomonadales commensals in the *tpst* mutant indicates a regulatory role of sulfated peptides rather than a confounding effect due to pleiotropic disorganization of the root MZ in this mutant. ANAC060 may also be involved in this interaction, although the evidence remains circumstantial. Genetic analysis of ANAC060 and other related ANAC family TFs is needed to disentangle the transcriptional regulatory machinery associated with and possibly responsible for Rhizobiales RGP activity.

Our results show that two independent lineages in Alpha- and Gammaproteobacteria classes employ independent mechanisms to promote host root growth, demonstrating that root-associated bacteria have likely acquired RGP capabilities more than once, independently. This, together with the wide prevalence of TPST-dependent RGP in extant Rhizobiales commensals, implies a benefit for bacteria in promoting host root growth, which has provided positive selective pressure on this trait. Whether this bacterial pathway actively aims to manipulate host growth or is passively utilized by hosts to alter their root growth remains to be addressed. A growing collection of microbial strains from a wide range of plant species, including those from an aquatic alga, *Chlamydomonas reinhardtii* (Durán et al., 2021), provides a useful tool to address this question. Moreover, the identification of bacterial genes responsible for RGP phenotype is crucial to genetically test the impact of this trait on plant and host fitness. We here provide the basis to better understand the molecular dialog between *A. thaliana* roots and Rhizobiales commensals to genetically address the relevance of microbiota-influenced regulation of plant growth and development. It is intriguing to test whether observed TPST-dependent manipulation of host root development has contributed to the subsequent acquisition of the ability to induce nodule organogenesis in the legume host.

## Methods

### Plant materials and growth conditions

We used *A. thaliana* wild-type accession Col-0 (CS60000) and commensal bacterial strains isolated from *A. thaliana* grown in natural soils, as described previously (Bai et al., 2015; Garrido-Oter et al., 2018). Plants were grown under short-day conditions (10 h under light at 21 °C and 14 h under dark at 19°C) in a square plate (120×120 mm^2^) placed vertically within a climate-controlled light cabinet (MLR-352, PHCbi).

### Bacterial culture

We used TY media (5 g/L tryptone, 3 g/L yeast extract, 10 mM CaCl_2_) to grow R129_E, R13_A, Root491, and Root142, and half strength of Tryptic Soy Broth media (15 g/L TSB; Sigma-Aldrich) to grow Root1497 and Root700. Bacterial cells were stored in glycerol stocks (16% v/v) at −80°C and recovered on TY supplemented with 1.8% agar, or 50% TSB supplemented with 1.5% agar, and cultivated at 28°C under dark conditions.

### Inoculation of plants with bacteria, root length quantification, and bacterial abundance quantification

Inoculation of plants with Alphaproteobacteria was performed as described previously (Garrido-Oter et al., 2018). Briefly, we cultivated bacteria in liquid TY or 50% TSB media for 2 days at 28°C with 180 rpm agitation and mixed bacterial cells into half-strength of Murashige and Skoog media (750 mM MgSO_4_, 625 mM KH_2_PO_4_, 10.3 mM NH_4_NO_3_, 9.4 mM KNO_3_, 1.5 mM CaCl_2_, 55 nM CoCl_2_, 53 nM CuCl_2_, 50 mM H_3_BO_3_, 2.5 mM KI, 50 mM MnCl_2_, 520 nM Na_2_MoO_4_ 15 mM ZnCl_2_, 75 mM Fe-EDTA, 500 mM MES-KOH pH 5.5) supplemented with 1% Bacto Agar (Difco). Bacterial titer (OD_600_) was adjusted to 0.00005 by mixing 5 μl of bacterial suspension (OD_600_ = 0.5 in 10 mM MgCl_2_) into 50 ml of MS media pre-cooled to around 40-50°C. *A. thaliana* seeds were directly placed onto bacterium-containing media or precultured on half-strength of Murashige and Skoog media with vitamin (Duchefa) supplemented with 1% Bacto Agar for 1 week and before seedlings were transferred to the bacterium-containing plates. The same media was supplemented with 1 μM flg22 (QRLSTGSRINSAKDDAAGLQIA; EZBiolab), 100 nM RGF1 [D-Y(SO3H)-SNPGHHP-Hyp-RHN; Beijing Scilight Biotechnology], or dRGF1 [DYSNPGHHP-Hyp-RHN] when necessary. Plates were then recorded by a scanner for root growth quantification using Fiji (Schindelin et al., 2012). Obtained data were square root transformed and fitted to a linear mixed model using biological and technical replicates as random factors, using the lmer function of the *lme4* R package. Fitted data was diagnosed to ensure the normal distribution of residuals and used for Student’s *t* test followed by correction for multiple comparison based on the Benjamini-Hochberg method. For bacterial biomass quantification, roots were thoroughly washed twice in sterile water to avoid residual agar pieces and frozen in a 2 ml screw-cap tubes with 1-mm zirconia beads in liquid nitrogen. Quantitative PCR was performed using the genomic DNA extracted from roots by DNeasy Plant Mini Kit (QIAGEN) and primer sets specific to *16S* rRNA gene sequences of *A. thaliana* (5’-CAGGCGGTGGAAACTACCAAG-3’ and 5’-TACAGCACTGCACGGGTCGAT-3’) and R129_E (5’-CGAGCTAATCTCCAAAAGCCATC-3’ and 5’-TGACCCTACCGTGGTTAGCTG-3’) with iQ SYBR Green Supermix (BIO-RAD), as described previously (Garrido-Oter et al., 2018).

### RNA extraction and sequencing

For transcriptomic analysis of roots inoculated with Alphaproteobacteria commensals, roots inoculated with bacteria for 12 days, as described above, were harvested and immediately frozen in liquid nitrogen. Roots from three plates (approximately 25 plants) were pooled into one sample. Three full-factorial biological replicates from three fully independent experiments were prepared for each condition. Roots were then homogenized with 1-mm zirconia beads (Carl Roth) and Precellys24 (6,300 rpm for 30 sec, 2 times with 15-sec intervals; Bertin), and RNA was extracted and purified with the RNeasy Plant Mini kit (QIAGEN). Quality control, library preparation, and sequencing (on the Illumina HiSeq 3000 platform) were conducted at the Max Planck Genome Centre (MPGC), Cologne. Approximately 20,000,000 reads (150-bp single-end) were obtained per sample.

For transcriptomic analysis of roots inoculated with R129_E in the presence of flg22, plants were inoculated as described above. Plants from one plate were pooled into one sample (approximately 10 plants) and three independent samples were obtained per condition. Two fully independent experiments were conducted and six samples per condition were obtained. Quality control, library preparation, and sequencing (on the Illumina HiSeq 3000 platform) were conducted at the Max Planck Genome Centre (MPGC), Cologne. Approximately 6,000,000 reads (150-bp single-end) were obtained per sample.

For transcriptomic analysis of roots inoculated with *B. phytofirmans* and *B. glumae,* Col-0 plants were grown on vertical agarose plates with Long Ashton nutrient solution adjusted for sulfate concentration of 750 μM (control) or 25 μM (low S) as described before (Koprivova et al., 2019). Total RNA was isolated from 18-day-old (14 dpi) roots and sequenced at the Genomics & Transcriptomics Laboratory (GTL) at Heinrich Heine University, Düsseldorf. Approximately 40,000,000 reads (150-bp single-end) were obtained per sample.

### RNA-seq analysis

Obtained reads, as well as reads downloaded from the Sequence Read Archive (SRA) database, were preprocessed by fastp (Chen et al., 2018) with default parameters and aligned to *A. thaliana* Col-0 genome by HISAT2 (Kim et al., 2019) with default parameters based on the latest gene annotation (Cheng et al., 2017). Reads per gene were counted by featureCounts (Liao et al., 2014) with default parameters. Subsequent statistical analyses were performed using R software (https://www.r-project.org/) unless described otherwise. Differential expression analysis was performed using the *edgeR* (Robinson et al., 2010) package. Library size was normalized by the weighted trimmed mean of M-values (TMM) method, and normalized read counts were fitted to a generalized linear model (GLM) with a negative binomial distribution to identify significantly differentially expressed genes (DEGs). Zero-centered expression values were calculated based on log_2_ counts per million (log2cpm) by subtracting the mean for each gene across samples. For comparison between datasets, TMM normalization, GLM fitting, and zero-centered expression values were calculated within each dataset. Zero-centered expression values were used to compute *k*-means clustering, where *k* was decided based on Bayesian Information Criterion (BIC). The similarity between samples was inferred by computing Pearson’s Correlation Coefficients (PCC) based on zero-centered expression values of DEGs (n = 1,439). Distance of PCCs to 1 was used as dissimilarity indices to perform principal coordinates analysis (PCoA) by cmdscale function. GO enrichment analysis was performed by Metascape online software (Zhou et al., 2019). Transcription binding motif enrichment analysis was performed using the ame function implemented in the MEME Suite (McLeay and Bailey, 2010). Gene regulatory network inference was performed using the *GENIE3* package (Huynh-Thu et al., 2010) with default parameters. We compared responses to different microbes by computing logFC within each dataset and use it to compute cosine similarity by the cosine function in the packge *lsa* and performed *k*-means clustering, whose *k* was determined based on the Bayesian Information Criterion. Distance of cosine similarities to 1 was used as dissimilarity indices to perform PCoA and CAP. Permutational analysis of variance (PERMANOVA) and pairwise PERMANOVA were performed by the adonis2 and pairwise.perm.manova functions of the *vegan* and *RVAideMemoire* packages, respectively. All plots were visualized by Metascape or by the *ggplot2, ggtree, patchwork,* and *igraph* packages in R.

### Single-cell RNA-seq analysis

T-distributed Stochastic Neighbor Embedding (t-SNE) and representation of clusterwise expression levels as a violin plot were created by The Plant Single Cell RNA-Sequencing Database (https://www.zmbp-resources.uni-tuebingen.de/timmermans/plant-single-cell-browser/). To perform co-expression analysis, read count data of 27,629 genes from 4,727 cells was retrieved from the NCBI Sequence Read Archive (SRA) under the accession number GSE123818, followed by global-scaling normalization by the NormalizeData function in the *Seurat* package (Hao et al., 2020). We then filtered for 22,017 highly variable genes and their expression values were scaled and centered by the FindVariableGenes and ScaleData functions. Normalized and scaled expression values were used to compute mutual ranks as a coexpression index for all pairs of genes as described previously (Obayashi and Kinoshita, 2009). The top 100 genes co-expressed with *ANAC060* were selected based on the mutual ranks and their GO enrichment was analyzed by Metascape.

### EdU staining and confocal microscopy

For EdU staining, roots of seed-inoculated plants at 21 dpi were incubated with EdU for two hours directly on agar plates, followed by fixation with 2% formaldehyde, 0.3% glutaraldehyde, 0.1% Triton X-100, 10 mM EGTA, and 5 mM MgSO_4_ in 50 mM sodium phosphate buffer (pH 7.2) for 1 hour, washing with phospho-buffered saline, and staining with Alexa488-azide using the Click-iT EdU Cell Proliferation Kit (Thermo Fisher). Tips (1-2 cm) of stained or non-stained roots were excised on agar plates and mounted onto a slide glass with Milli-Q water or 10 μg/ml propidium iodide (PI). PI and GFP fluorescence were observed by a SP8 FALCON-DIVE confocal microscope (Leica) using a 2-photon laser for excitation at 880 nm for GFP and 1,045 nm for PI and emission at 498-521 for GFP and 606-630 nm for PI, or by a LSM780 confocal microscope (Zeiss) using a 25-mW Argon ion laser for excitation at 488 nm and emission at 490-534 nm for GFP and Alexa488. Acquired images were processed and adjusted (equally within each experiment) by Fiji (Schindelin et al., 2012).

### Data and code availability

Raw sequencing reads obtained in this study are deposited to the European Nucleotide Archive (ENA) under the accession numbers of PRJEB45043 (Alphaproteobacteria), PRJEB45044 (R129_E with flg22), and PRJEB45045 *(Burkholderia).* The scripts for the computational analyses described in this study are available at https://github.com/rtnakano1984/RNAseq_Alphaproteobacteria and https://github.com/rtnakano1984/RNAseq_compare to ensure replicability and reproducibility of these results.

## Acknowledgments

We would like to thank Paul Schulze-Lefert at the Max Planck Institute for Plant Breeding Research for providing infrastructures and helpful advice for this study, Jiri Friml at the Institute of Science and Technology Austria, Niko Geldner at the University of Lausanne, Woei-Jiun Guo at the National Cheng Kung University, Yoshikatsu Matsubayashi and Hidefumi Shinohara at the Nagoya University, Birgit Kemmerling at the University of Tübingen, Jia Li at the Lanzhou University, Margret Sauter at the University of Kiel, and Bruno Müller at the University of Zurich for providing *A. thaliana* mutant and transgenic seeds, Bruno Hüttel and Mark Holstein at the MPGC and Patrick Petzsch and Karl-Erich Köhler at the GTL for sequencing, and Christina Phillip and Petra Köchner for their technical support. We also would like to thank Tatsuya Nobori, Kenichi Tsuda, Stephane Hacquard, and Rozina Kardakaris for their critical reading and help in editing the manuscript. This work was supported by the Deutsche Forschungsgemeinschaft (DFG, German Research Foundation) under the Priority Programme “2125 Deconstruction and Reconstruction of Plant Microbiota (DECRyPT)” funded to RTN (402201269) and SK (401836049). This work was also supported by the Cluster of Excellence on Plant Sciences (CEPLAS) funded by DFG to TG, SK, and GWK. The authors declare no conflicts of interest. In memory of Alina Kuroczik, who provided technical support for this study.

## Author contribution

RTN conceived and designed the research and performed experiments. JH, TG, and AK performed experiments partly under supervision of RTN and SK. PS, MNC and GWK contributed to the RNAseq data analysis. JH and RTN analyzed and interpreted the data. RTN wrote the paper with inputs from the other authors.

---

**Supplemental Figure 1. Qualitative comparison of DEGs for Alphaproteobacteria inoculations**

Upset diagrams showing the number of DEGs specific to or shared (intersect) between different inoculations. Up-regulated and down-regulated DEGs were separately analyzed.

**Supplemental Figure 2. Similarity in gene contents between GO categories enriched in Alphaproteobacteria DEGs based on *k*-means clustering.**

For each cluster, significant enriched GO categories are shown in rows. Genes in the respective clusters with at least one significantly enriched GO category are shown in columns. Assignment of a gene to a given GO category is indicated by black filling color. GO categories with highly similar gene contents are manually marked and representative terms are indicated. Bold R and B in the labels indicate “response to” and “biosynthesis of”, respectively.

**Supplemental Figure 3. WRKY54 and ANAC060 are associated with responses characteristic to sister lineage and Rhizobiales isolates, respectively.**

The same GRN as in Figure 2B, colored by k-means clusters (A), by logFC upon Alphaproteobacteria commensals (B; the full view of Figure 3C), and for Rhizobiales-specific DEGs (C; the full view of Figure 3B). Module 1 and 2 contain genes from Clusters 1, 8, and 9 and Clusters 3, 4, and 6, respectively, and Rhizobiales DEGs were exclusively found in the module 2. Part of the network presented in Figures 4B and 4C are indicated by a square.

**Supplemental Figure 4. Robust centrality of WRKY54 and ANAC060 irrespective of the lists of potential regulators.**

GENIE3-based inference of gene regulatory network using all differentially expressed transcription factors as potential regulators, colored for transcription factor families (A), by k-means clusters, corresponding to Figure 1B (B), for Rhizobiales-specific DEGs, corresponding to Figure 3A (C) or by logFC (D). Modules are arbitrarily defined. (E) Centrality indices of potential regulators in the inferred GRN.

**Supplemental Figure 5. Non-pathogenic microbial inoculation triggered similar transcriptional responses in the host roots.**

(A) A heatmap showing logFC of genes differentially expressed at least in one of the analyzed treatments (n = 13,889), aligned according to *k*-means clustering (*k* = 21, based on Bayesian information criterion). Assignment of genes to *k*-means cluster for Alphaproteobacteria dataset (Figure 1) are also shown above. (B) GO enrichment analysis of k-means clustering shown in (A), as in Figure 1F. (C) A heatmap showing logFC of GST genes, aligned according to their phylogeny. Shaded by blue are the genes specifically acquired within the family Brassicaceae (Pisléska-Bednarek et al., 2018).

**Supplemental Figure 6. ANAC060 is expressed in the cell population within the root meristematic zone, where PLETHORA TFs and RGF peptides are also expressed to control meristematic mitotic activity.**

(A) T-distributed stochastic neighbor embedding (t-SNE) of single cell RNAseq datasets reported by Denyer and Me et al. (2019), where colors correspond to the normalized expression levels of genes indicated above. (B) Cluster assignment of ANAC060, PLT1, PLT2, and PLT4, where the cluster 11 corresponds to the meristematic cell population. t-SNE and violin plots were created by The Plant Single Cell RNA-Sequencing Database. (C) GO enrichment analysis of 100 genes best co-expressed with ANAC060 in the single cell RNAseq datasets, based on Pearson’s correlation-based Mutual Ranks. Data are visualized as in Figure 1F.

**Supplemental Figure 7. DNA synthesis and auxin and cytokinin accumulation in axenic and R129_E-inoculated roots.**

(A) Roots incubated with EdU for 2 hours were staining with Alexa488 after seeds were inoculated with R129_E for 21 days under short-day conditions. EdU/Alexa488 signals in green was merged with autofluorescence signals in magenta. (B) Roots expressing DR5:ER-GFP were imaged after seeds were inoculated with R129_E for 8 days under short-day condition. (C) Roots expressing TSCn:ER-GFP were imaged after 7-day-old seedlings were inoculated with R129_E for 10 days under short-day conditions. Bars, 50 μm.

**Supplemental Figure 8. RGF and CIF peptide pathways were not essential for R129_E RGP activity.**

(A) R129_E promoted primary root growth of Col-0, *rgf123*, and *sgn3* plants after 7-day-old seedlings were inoculated for 14 days under short-day conditions. (B) External application of 100 nM RGF1 or 100 nM desulfated RGF1 did not affect the ability of R129_E to promote root growth of Col-0 and did not rescue the lack of RGP in *tpst* mutant roots. Letters indicate statistical significance (α = 0.05) corresponding to Student’s *t* test after fitting to a linear mixed model with biological and technical replicates as random factors, corrected for multiple comparison by the Benjamini-Hochberg method. (C) Roots of *tpst* mutant plants were stained with 10 μg/mL propidium iodide after 7-day-old seedlings were inoculated with R129_E for 14 days in the presence or absence of 100 nM RGF1. Arrowheads indicate the site of transition from cell division to cell elongation. Bars, 50 μm.

**Supplemental Table 1. RNAseq datasets analyzed in this study.**

**Supplemental Table 2. Pairwise PERMANOVA of logFC by lifestyles**

