## Supplemental for "Rhizobiales commensal bacteria promote *Arabidopsis thaliana* root growth via host sulfated peptide pathway"

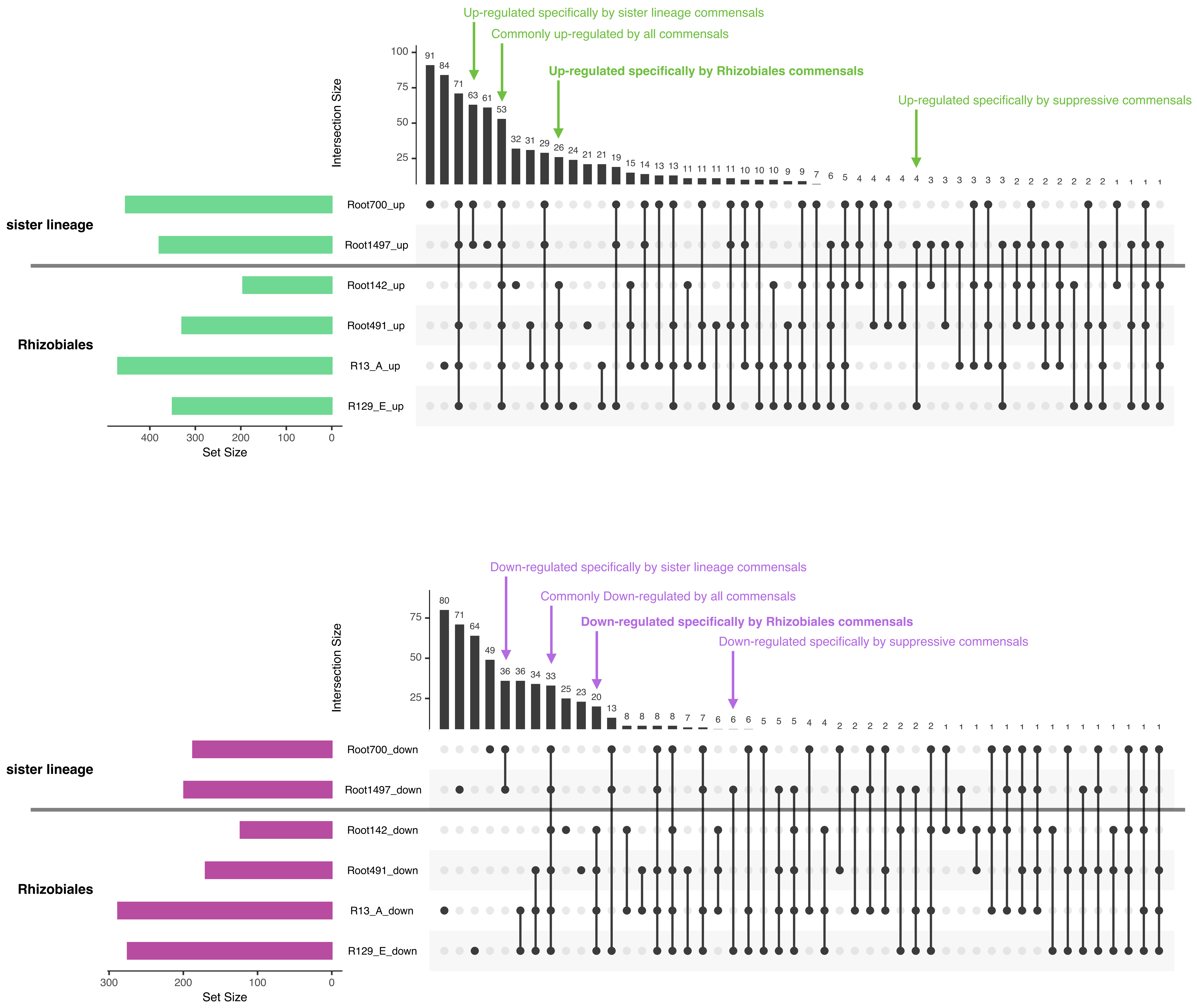


**Supplemental Figure 1. Qualitative comparison of DEGs for Alphaproteobacteria inoculations**
Upset diagrams showing the number of DEGs specific to or shared (intersect) between different inoculations. Up-regulated and down-regulated DEGs were separately analyzed.

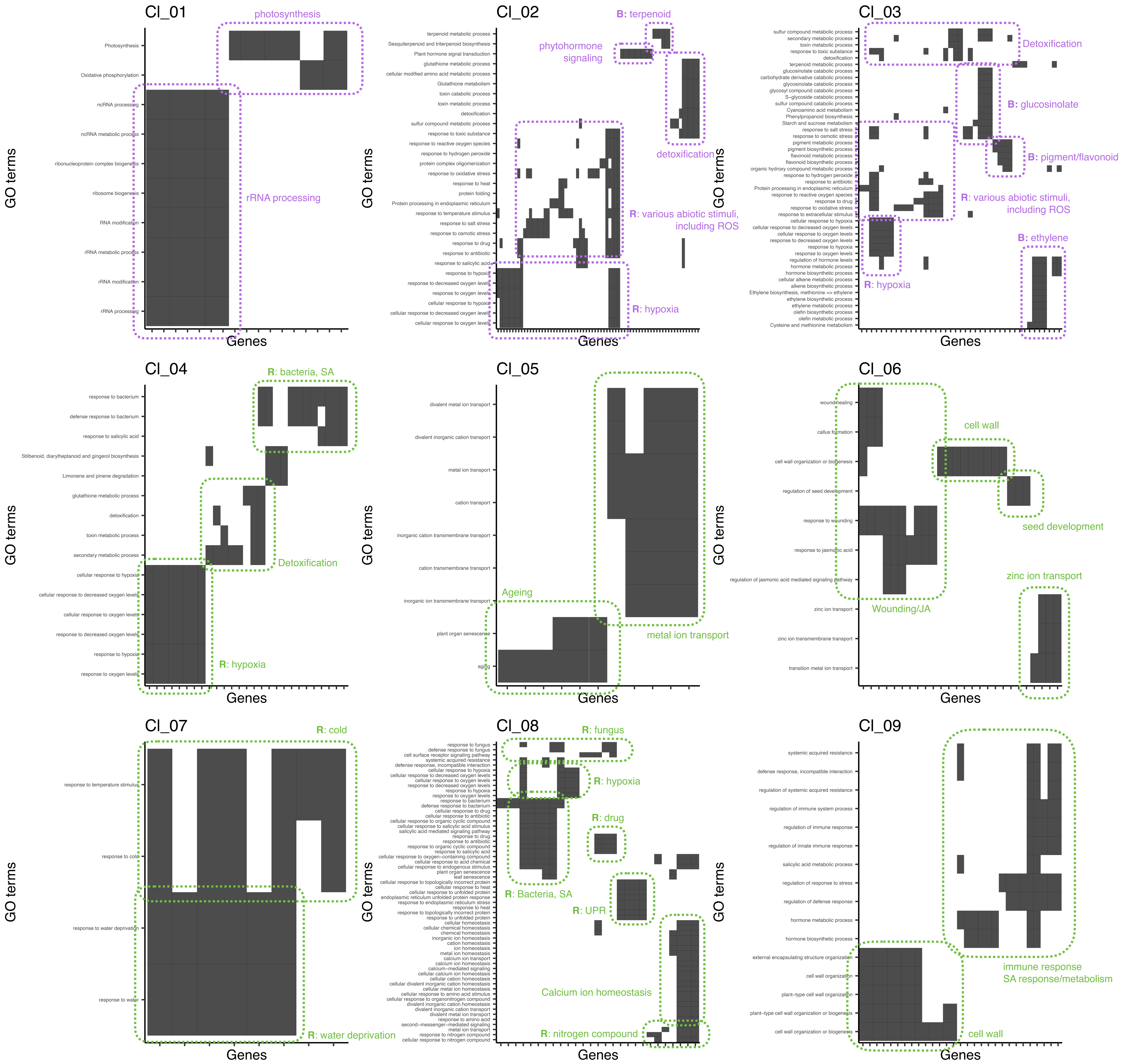


**Supplemental Figure 2. Similarity in gene contents between GO categories enriched in Alphaproteobacteria DEGs based on *k*-means clustering.**

For each cluster, significant enriched GO categories are shown in rows. Genes in the respective clusters with at least one significantly enriched GO category are shown in columns. Assignment of a gene to a given GO category is indicated by black filling color. GO categories with highly similar gene contents are manually marked and representative terms are indicated. Bold R and B in the labels indicate "response to" and "biosynthesis of", respectively.

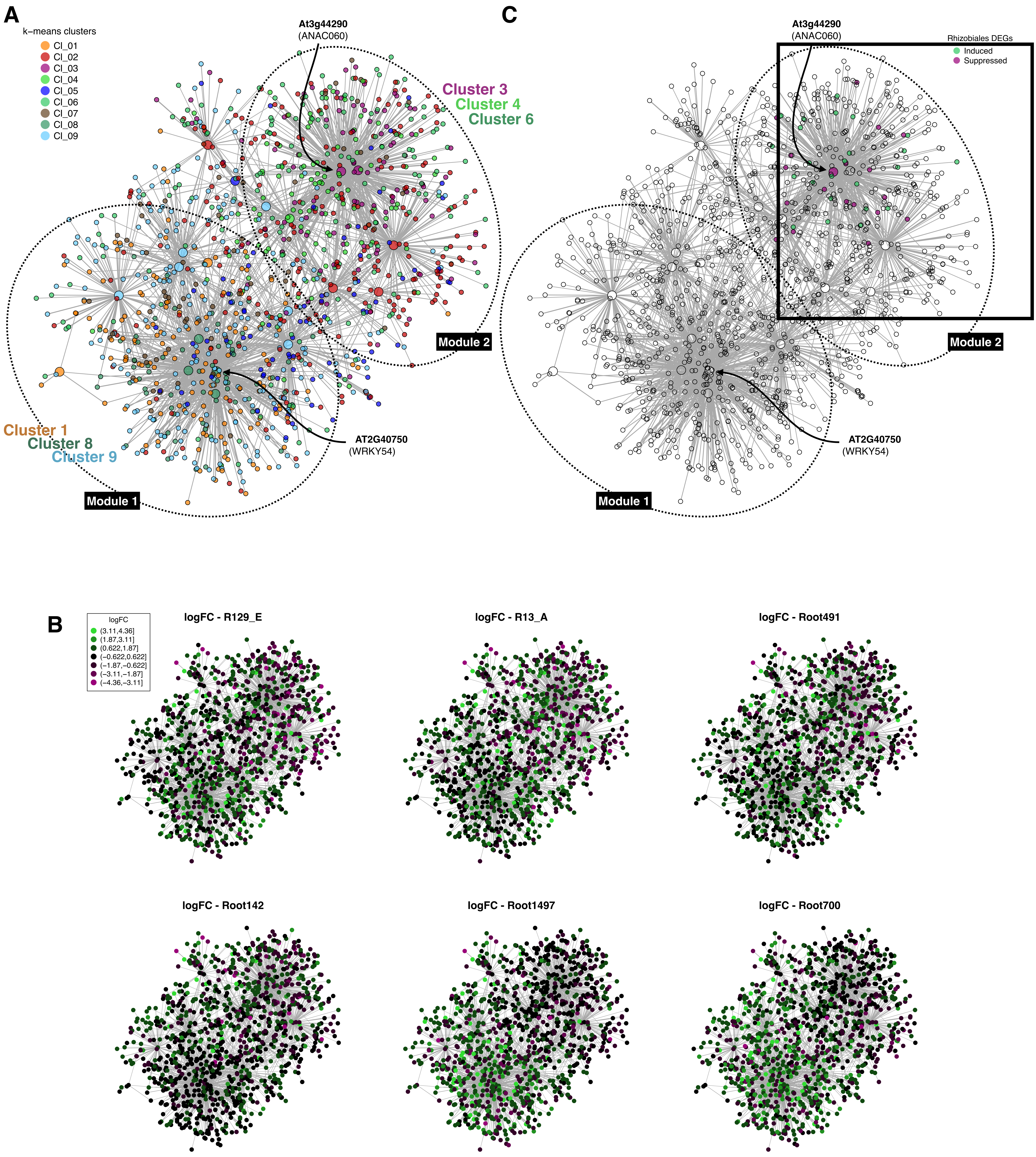


**Supplemental Figure 3. WRKY54 and ANAC060 are associated with responses characteristic to sister lineage and Rhizobiales isolates, respectively.**

The same GRN as in Figure 2B, colored by k-means clusters (A), by logFC upon Alphaproteobacteria commensals (B; the full view of Figure 3C), and for Rhizobiales-specific DEGs (C; the full view of Figure 3B). Module 1 and 2 contain genes from Clusters 1, 8, and 9 and Clusters 3, 4, and 6, respectively, and Rhizobiales DEGs were exclusively found in the module 2. Part of the network presented in Figures 4B and 4C are indicated by a square.

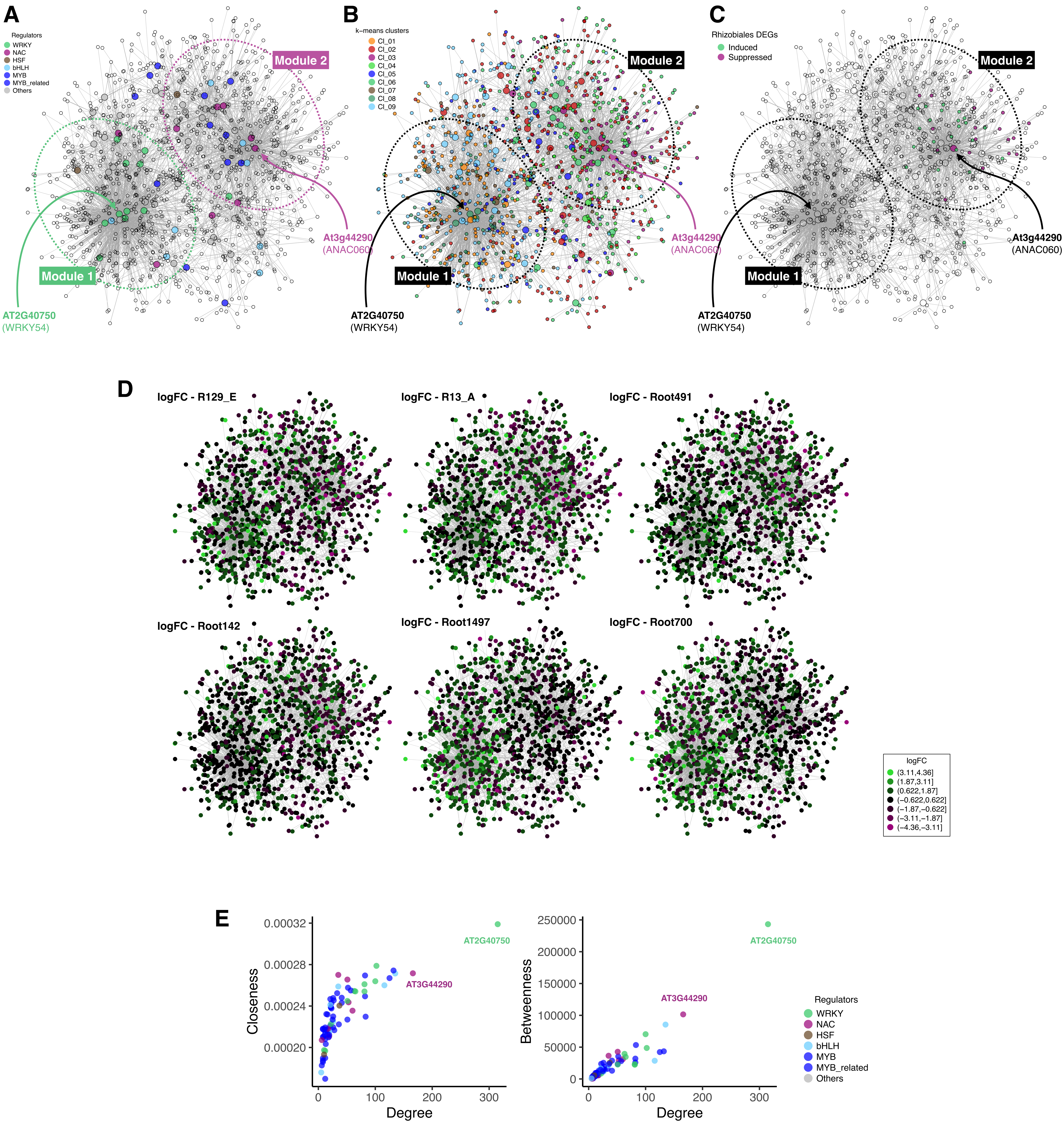


**Supplemental Figure 4. Robust centrality of WRKY54 and ANAC060 irrespective of the lists of potential regulators.** GENIE3-based inference of gene regulatory network using all differentially expressed transcription factors as potential regulators, colored for transcription factor families (A), by k-means clusters, corresponding to Figure 1B (B), for Rhizobiales-specific DEGs, corresponding to Figure 3A (C) or by logFC (D). Modules are arbitrarily defined. (E) Centrality indices of potential regulators in the inferred GRN.

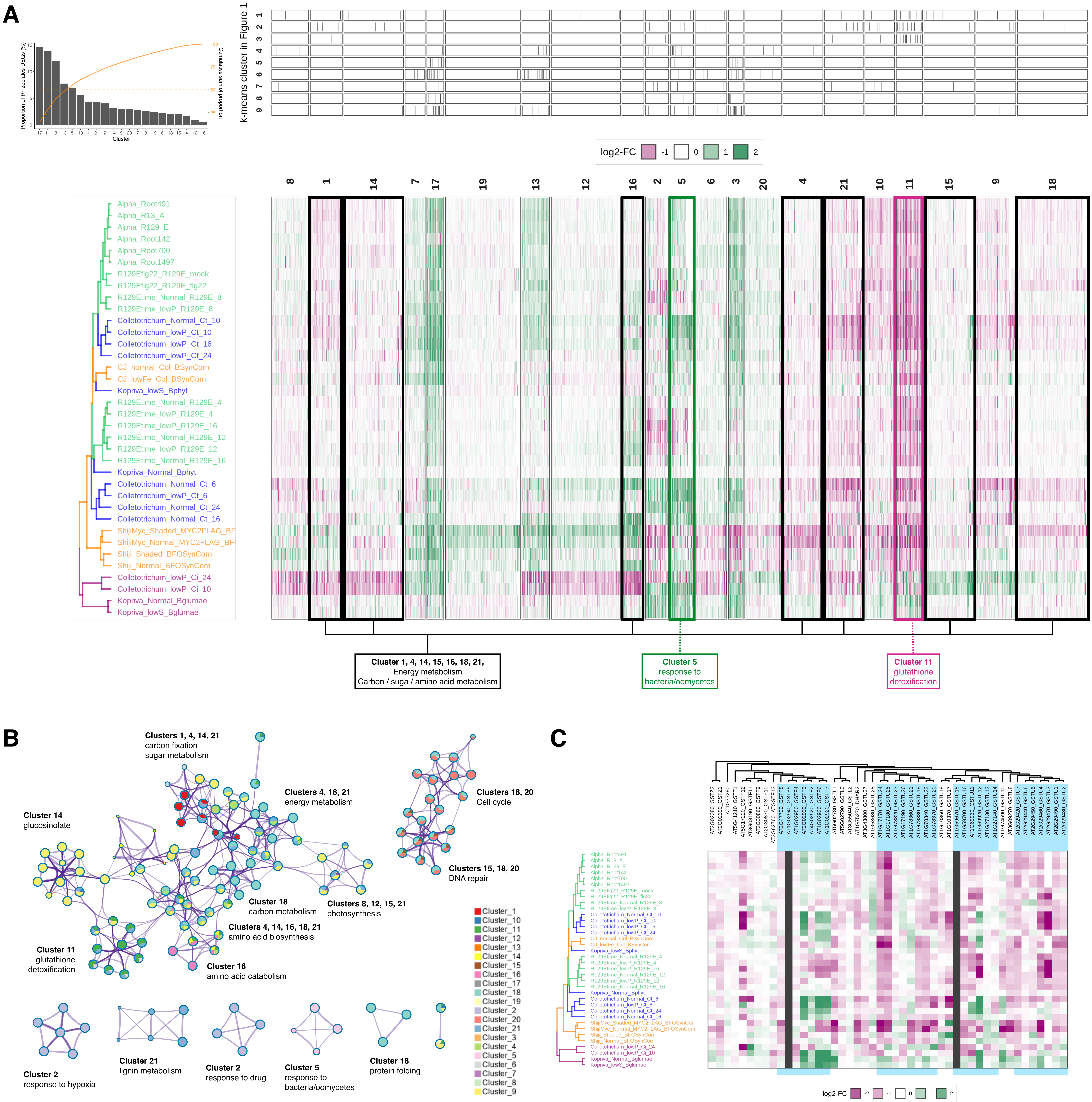


**Supplemental Figure 5. Non-pathogenic microbial inoculation triggered similar transcriptional responses in the host roots.** (A) A heatmap showing logFC of genes differentially expressed at least in one of the analyzed treatments (n = 13,889), aligned according to k-means clustering (k = 21, based on Bayesian information criterion). Assignment of genes to *k*-means cluster for Alphaproteobacteria dataset (Figure 1) are also shown above. (B) GO enrichment analysis of k-means clustering shown in (A), as in Figure 1F. (C) A heatmap showing logFC of GST genes, aligned according to their phylogeny. Shaded by blue are the genes specifically acquired within the family Brassicaceae (Pisléska-Bednarek et al., 2018).

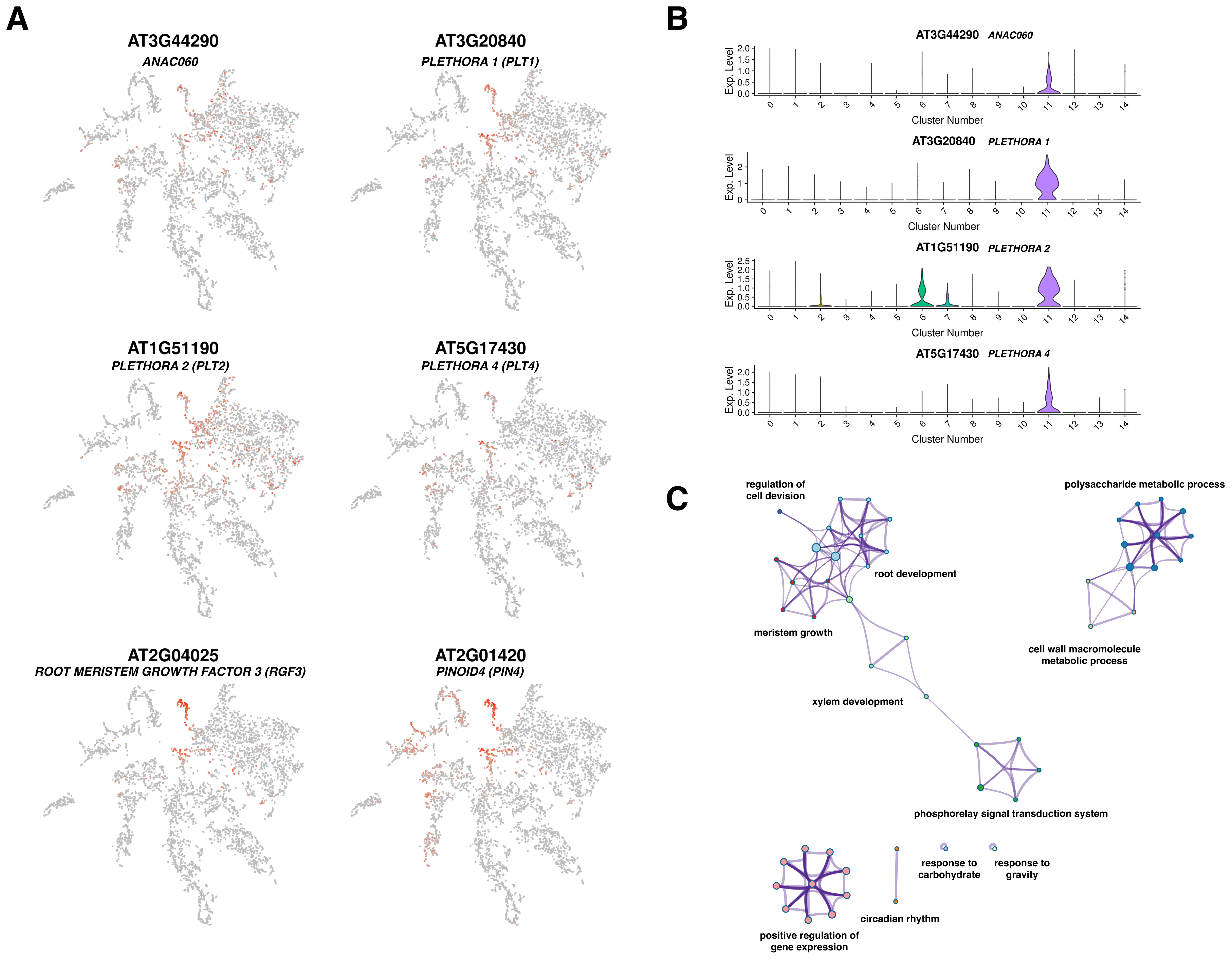


**Supplemental Figure 6. ANAC060 is expressed in the cell population within the root meristematic zone, where PLETHORA TFs and RGF peptides are also expressed to control meristematic mitotic activity.**
(A) T-distributed stochastic neighbor embedding (t-SNE) of single cell RNAseq datasets reported by Denyer and Me et al. (2019), where colors correspond to the normalized expression levels of genes indicated above. (B) Cluster assignment of ANAC060, PLT1, PLT2, and PLT4, where the cluster 11 corresponds to the meristematic cell population. t-SNE and violin plots were created by The Plant Single Cell RNA-Sequencing Database. (C) GO enrichment analysis of 100 genes best co-expressed with ANAC060 in the single cell RNAseq datasets, based on Pearson's correlation-based Mutual Ranks. Data are visualized as in Figure 1F.

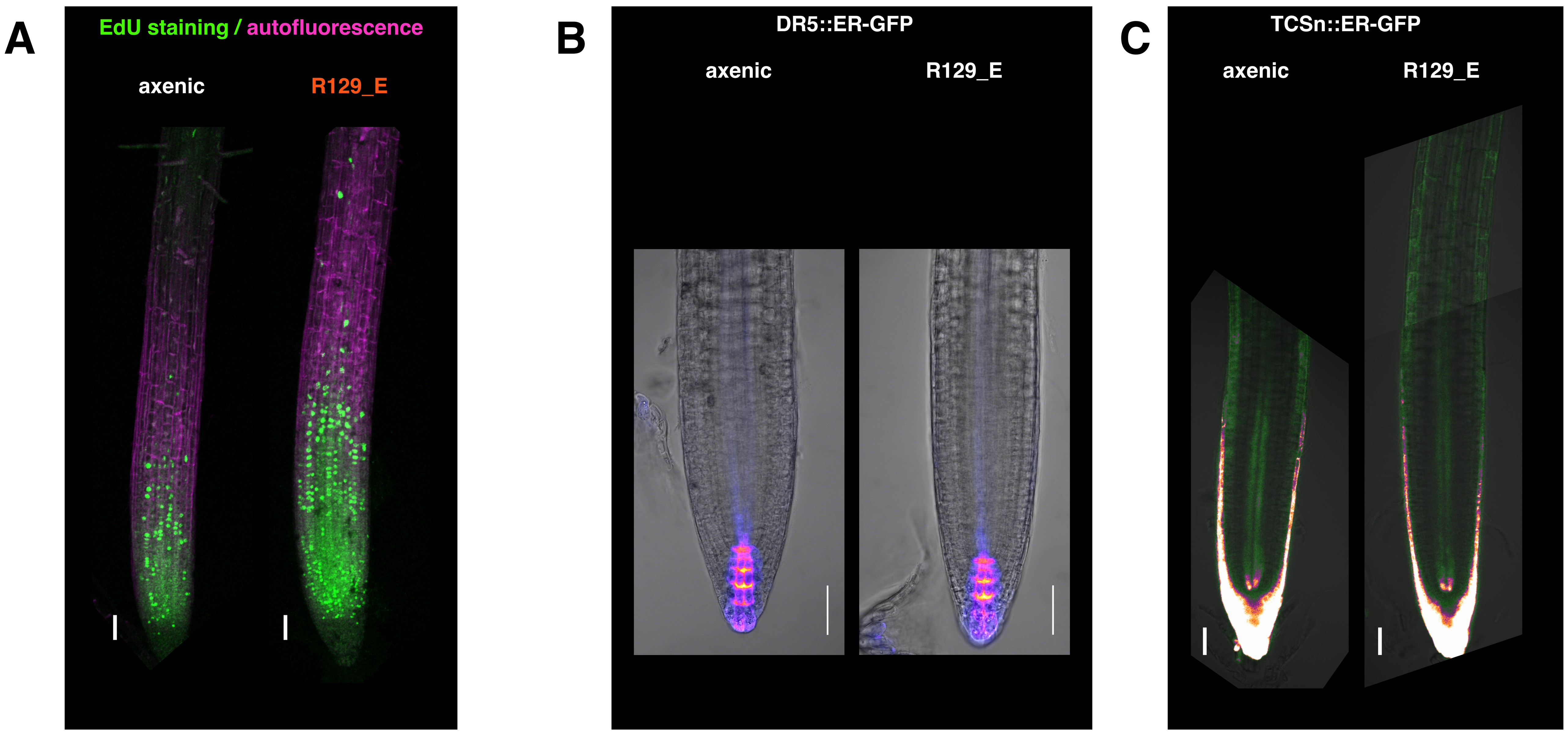


**Supplemental Figure 7. DNA synthesis and auxin and cytokinin accumulation in axenic and R129_E-inoculated roots.**
(A) Roots incubated with EdU for 2 hours were staining with Alexa488 after seeds were inoculated with R129_E for 21 days under short-day conditions. EdU/Alexa488 signals in green was merged with autofluorescence signals in magenta. (B) Roots expressing DR5::ER-GFP were imaged after seeds were inoculated with R129_E for 8 days under short-day condition. (C) Roots expressing TSCn::ER-GFP were imaged after 7-day-old seedlings were inoculated with R129_E for 10 days under short-day conditions. Bars, 50 μm.

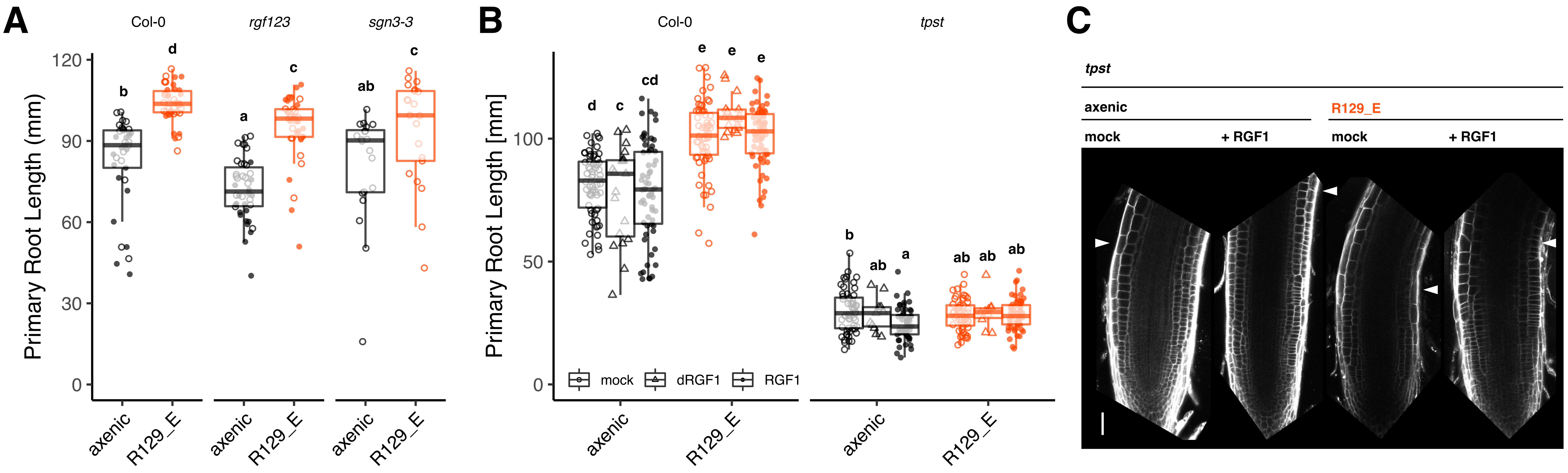


**Supplemental Figure 8. RGF and CIF peptide pathways were not essential for R129_E RGP activity.**
(A) R129_E promoted primary root growth of Col-0, *rgf123*, and *sgn3* plants after 7-day-old seedlings were inoculated for 14 days under short-day conditions. (B) External application of 100 nM RGF1 or 100 nM desulfated RGF1 did not affect the ability of R129_E to promote root growth of Col-0 and did not rescue the lack of RGP in *tpst* mutant roots. Letters indicate statistical significance (α = 0.05) corresponding to Student's *t* test after fitting to a linear mixed model with biological and technical replicates as random factors, corrected for multiple comparison by the Benjamini-Hochberg method. (C) Roots of *tpst* mutant plants were stained with 10 μg/mL propidium iodide after 7-day-old seedsling were inoculated with R129_E for 14 days in the presence or absence of 100 nM RGF1. Arrowheads indicate the site of transition from cell division to cell elongation. Bars, 50 μm.

| Inoculant | Inoculant lifestyle | Inoculant kingdom | Additional treatment | Inoculation system | Nutrient | Light | Genotype | Time point (dpi) | Reference |
|---|---|---|---|---|---|---|---|---|---|
| R129_E<br>R13_A<br>Root491<br>Root142<br>Root1497<br>Root700 | Commensal | B | n.a. | Agar | Normal 1/2 MS | Normal, short-day<br>(10 h light / 14 h dark) | Col-0 | 12 | This study |
| B-SynCom | SynCom | B | n.a. | Agar | 1/2 MS with avFe (Fe-EDTA)<br>1/2 MS with unavFe ($FeCl_3$) | Normal, short-day<br>(10 h light / 14 h dark) | Col-0 | 7 | 1 |
| *C. tofieldiae*<br>*C. incanum* | Beneficial<br>Pathogenic | F | n.a. | Agar | 1/2 MS with high phosphate (625 µM)<br>1/2 MS with low phosphate (50 µM) | Normal, short-day<br>(10 h light / 14 h dark) | Col-0 | 6, 10, 16, 24 | 2,3 |
| *B. phytofirmens*<br>*B. glumae* | Beneficial<br>Pathogenic | B | n.a. | Agar | 1/2 MS with high sulfate (750 µM)<br>1/2 MS with low sulfate (25 µM) | Normal, long-day<br>(16 h light / 8 h dark) | Col-0 | 14 | This study |
| R129_E | Commensal | B | flg22 | Agar | Normal 1/2 MS | Normal, short-day<br>(10 h light / 14 h dark) | Col-0 | 12 | This study |
| R129_E | Commensal | B | n.a. | Agar | 1/2 MS with high phosphate (625 µM)<br>1/2 MS with low phosphate (50 µM) | Normal, short-day<br>(10 h light / 14 h dark) | Col-0 | 4, 8, 12, 16 | 4 |
| BFO-SynCom | SynCom | BFO | n.a. | FlowPot | Peat-based sterilized soil matrix | Normal or Shaded,<br>short-day<br>(10 h light / 14 h dark) | Col-0 | 35 | 5 |
| BFO-SynCom | SynCom | BFO | n.a. | FlowPot | Peat-based sterilized soil matrix | Normal or Shaded,<br>short-day<br>(10 h light / 14 h dark) | MYC2-FLAG in<br>*myc2-3* | 35 | 5 |


**Supplemental Table 1. RNAseq datasets analyzed in this study.**

| Lifestyles | | $R^2$ | FDR-corrected *P* value |
|---|---|---|---|
| pathogenic | SynCom | 0.440 | 0.0100 |
| pathogenic | commensal | 0.344 | 0.0015 |
| pathogenic | beneficial | 0.263 | 0.0015 |
| SynCom | beneficial | 0.255 | 0.0024 |
| SynCom | commensal | 0.229 | 0.0015 |
| commensal | beneficial | 0.201 | 0.0015 |

**Supplemental Table 2. Pairwise PERMANOVA of logFC by lifestyles**
